# An essential, kinetoplastid-specific GDP-Fuc: β-*D*-Gal α-1,2-fucosyltransferase is located in the mitochondrion of *Trypanosoma brucei*

**DOI:** 10.1101/726117

**Authors:** Giulia Bandini, Sebastian Damerow, Maria Lucia Sampaio Güther, Hongjie Guo, Angela Mehlert, Stephen M. Beverley, Michael A. J. Ferguson

## Abstract

Fucose is a common component of eukaryotic cell-surface glycoconjugates, generally added by Golgi-resident fucosyltransferases. Whereas fucosylated glycoconjugates are rare in kinetoplastids, the biosynthesis of the nucleotide sugar GDP-Fuc has been shown to be essential in *Trypanosoma brucei.* Here we show that the single identifiable *T. brucei* fucosyltransferase (TbFUT1) is a GDP-Fuc: β-D-galactose α-1,2-fucosyltransferase with an apparent preference for a Galβ1,3GlcNAcβ1-O-R acceptor motif. Conditional null mutants of *TbFUT1* demonstrated that it is essential for both the mammalian-infective bloodstream form and the insect vector-dwelling procyclic form. Unexpectedly, TbFUT1 was localized in the mitochondrion of *T. brucei* and found to be required for mitochondrial function in bloodstream form trypanosomes. Finally, the *TbFUT1* gene was able to complement a *Leishmania major* mutant lacking the homologous fucosyltransferase gene (Guo et al., 2021). Together these results suggest that kinetoplastids possess an unusual, conserved and essential mitochondrial fucosyltransferase activity that may have therapeutic potential across trypanosomatids.

## INTRODUCTION

The protozoan parasites of the *Trypanosoma brucei* group are the causative agents of human and animal African trypanosomiasis. Bloodstream form *T. brucei* is ingested by the tsetse fly vector and differentiates into procyclic form parasites to colonize the tsetse midgut. To then infect a new mammalian host, *T. brucei* undergoes a series of differentiations that allows it to colonize the fly salivary gland and to be transferred to a new host during a subsequent blood meal (Matthews, 2005).

The surface coat of the bloodstream form is characterized by the GPI-anchored*, N*-glycosylated and occasionally *O*-glycosylated variant surface glycoprotein (VSG) (Cross, 1996; Mehlert et al., 1998; Pays, 1998; Pinger et al., 2018; Schwede and Carrington, 2010), while procyclic cells express a family of GPI-anchored proteins called procyclins (Richardson et al., 1988; Roditi et al., 1998; Treumann et al., 1997; Vassella et al., 2001), free glycoinositolphospholipids (Nagamune et al., 2004; Roper et al., 2005; Vassella et al., 2003) and a high molecular weight glycoconjugate complex (Güther et al., 2009). The importance of glycoproteins to parasite survival and infectivity has led to the investigation of enzymes for GPI anchor biosynthesis (Chang et al., 2002; Nagamune et al., 2000; Smith et al., 2004; Urbaniak et al., 2008, 2014) and nucleotide sugar biosynthesis (Bandini et al., 2012; Denton et al., 2010; Keuttel et al., 2012; Guther et al, 2021; Marino et al., 2010, 2011; Roper et al., 2002, 2005; Stokes et al., 2008; Shaw et al., 2003; Turnock et al., 2007; Urbaniak et al., 2006a, 2006b; 2013; Zmuda et al. 2019) as potential therapeutic targets.

Nucleotide sugars are used as glycosyl donors in many glycosylation reactions. GDP-Fucose (GDP-Fuc) was identified in the nucleotide sugar pools of *T. brucei*, *Trypanosoma cruzi* and *Leishmania major* (Turnock and Ferguson, 2007) and its biosynthesis is essential for parasite growth in procyclic and bloodstream form *T. brucei* (Turnock et al., 2007) and in *L. major* promastigotes (Guo et al., 2017). Interestingly, *T. brucei* and *L. major* use different pathways to synthesize GDP-Fuc. *T. brucei* utilizes the *de novo* pathway in which GDP-Fuc is synthesised from GDP-Mannose via GDP-mannose-4,6-dehydratase (GMD) and GDP-4-keto-6-deoxy-D-mannose epimerase/reductase (GMER) (Guther et al., 2021; Turnock et al., 2007; Turnock and Ferguson, 2007). Conversely, *L. major* has two related bifunctional D-arabinose/L-fucose kinase/pyrophosphorylase, AFKP80 and FKP40, that synthesise GDP-Fuc from free fucose (Guo et al., 2017). Despite the aforementioned essentialities for GDP-Fuc in *T. brucei* and *L. major*, the only structurally-defined fucose-containing oligosaccharide in trypanosomatids is the low-abundance Ser/Thr-phosphodiester-linked glycan on *T. cruzi* gp72, a glycoprotein that has been implicated in flagellar attachment (Allen et al., 2013; Cooper et al., 1993; Ferguson et al., 1983; Haynes et al., 1996).

Fucosyltransferases (FUTs) catalyse the transfer of fucose from GDP-Fuc to glycan and protein acceptors and are classified into two superfamilies (Coutinho et al., 2003; Lombard et al., 2013). One superfamily contains all α1,3/α1,4-FUTs (Carbohydrate Active EnZyme, CAZy, family GT10) and the other contains all α1,2-, α1,6- and protein *O*-fucosyltransferases (GT11, GT23, GT37, GT56, GT65, GT68 and GT74) (Martinez-Duncker, 2003). In eukaryotes, the vast majority of fucosyltransferases are type II transmembrane Golgi proteins (Breton et al., 1998), but two exceptions have been described: (i) PgtA, a cytoplasmic bifunctional β1,3-galactosyltransferase / α1,2-FUT found in *Dictyostelium discoideum* and *Toxoplasma gondii* (Rahman et al., 2016; Van Der Wel et al., 2002) that is part of an oxygen-sensitive glycosylation pathway that attaches a pentasaccharide to the Skp1-containing ubiquitin ligase complex (West et al., 2010); and (ii) SPINDLY, a protein *O*-fucosyltransferase that modifies nuclear proteins in *Arabidopsis thaliana* and *T. gondii* (Gas-Pascual et al., 2019; Zentella et al., 2017).

*T. brucei* and other kinetoplastids contain a single mitochondrion. In the bloodstream form of the parasite this organelle has a tubular structure, while in the procyclic form it is organized in a complex network with numerous cristae, reflecting the absence and presence, respectively, of oxidative phosphorylation (Matthews, 2005; Priest and Hajduk, 1994). The parasite mitochondrion is further characterized by a disc-shaped DNA network called the kinetoplast (Jensen and Englund, 2012) that is physically linked with the flagellum basal body (Ogbadoyi et al., 2003; Povelones, 2014).

While secretory pathway and nuclear/cytosolic glycosylation systems have been studied extensively, little is known about glycosylation within mitochondria. A glycoproteomic approach in yeast revealed several mitochondrial glycoproteins (Kung et al., 2009), but it was not determined whether these were imported from the secretory pathway or glycosylated within the mitochondria by as yet unknown glycosyltransferases. The only characterised example of a mitochondrial glycosyltransferase is the mitochondrial isoform of mammalian *O*-GlcNAc transferase (OGT). *O*-GlcNAcylation is a cycling modification, involved in signalling, in which OGT adds GlcNAc to Ser/Thr residues and *O*-GlcNAcase (OGA) removes it (Bond and Hanover, 2015). Studies have shown that both mitochondrial OGT (mOGT) and OGA are present and active in mammalian mitochondria and putative mitochondrial targets have been identified (Banerjee et al., 2015; Sacoman et al., 2017). Further, a mammalian mitochondrial UDP-GlcNAc transporter associated with mitochondrial *O*-GlcNAcylation has been described (Banerjee et al., 2015). However, orthologues of OGT and OGA genes are not present in kinetoplastids.

Here, we report on a gene (*TbFUT1*) encoding a mitochondrial α-1,2-fucosyltransferase protein (TbFUT1) in *T. brucei* that is essential to parasite survival. Similar results were obtained in the related trypanosomatid parasite *Leishmania major* (Guo et al. 2021), extending this unexpected finding across the trypanosomatid protozoans.

## RESULTS

### Identification, cloning and sequence analysis of TbFUT1

The CAZy database lists eight distinct FUT families (see Introduction) (Lombard et al., 2013). One or more sequences from each family were selected for BLASTp searches of the predicted proteins from the *T. brucei, T. cruzi* and *L. major* genomes (Table S1). Strikingly, only one putative fucosyltransferase gene (*TbFUT1*) was identified in the *T. brucei* genome (GeneDB ID: Tb927.9.3600) belonging to the GT11 family, which is comprised almost exclusively of α-1,2-FUTs (Coutinho et al., 2003; Zhang et al., 2010). Homologues of *TbFUT1* were also found in the *T. cruzi* and *L. major* genomes and, unlike *T. brucei*, *T. cruzi* and *L. major* also encode for GT10 FUT genes (Table S1). Finally, *Leishmania* spp. express a family of α-1,2-arabinopyranosyltransferases (SCA1/2/L, CAZy family GT79) that decorate phosphoglycan side chains and have been suggested to act as fucosyltransferases in presence of excess fucose (Guo *et al*., 2021).

The TbFUT1 predicted amino acid sequence shows relatively low sequence identity to previously characterized GT11 FUTs, *e.g*. *H. pylori* (26%) or human FUT2 (21%) (Kelly et al., 1995; Wang et al., 1999). Nevertheless, conserved motifs characteristic of this family can be clearly identified (Fig. 1*A*) (Li et al., 2008; Martinez-Duncker, 2003). Motif I (aa 153-159) is shared with α-1,6-FUTs and has been implicated in the binding of GDP-Fuc (Takahashi et al., 2000), whereas no clear functions have yet been assigned to motifs II, III and IV (aa 197-207, 265-273 and 13-18, respectively). Although TbFUT1 contains a putative transmembrane (TM) domain at its N-terminus (residues 6-28), as would be expected of a typical Golgi-localized fucosyltransferase (Breton et al., 1998), it overlaps with the fucosyltransferase motif IV, which normally occurs after the TM domain (Fig. 1*A*). Indeed, further analysis of the TbFUT1 predicted amino acid sequence using *PSort II (Horton and Nakai, 1997)*, *Target P* (Emanuelsson et al., 2000) and *Mitoprot* (Claros and Vincens, 1996) suggested mitochondrial localization and identified a putative mitochondrial targeting motif (M/L…RR) with RR at sequence positions 30 and 31. Conservation of this eukaryotic targeting motif has been previously shown for other parasite mitochondrial proteins (Krnáčová et al., 2012; Long et al., 2008). Additional BLASTp searches showed that there is generally a single TbFUT1 gene homologue in each kinetoplastid species, and a phylogram of FUT sequences indicates that TbFUT1 homologues form a distinct clade closest to bacterial α-1,2-fucosyltransferases (Fig. 1*B*).

**FIGURE 1.**
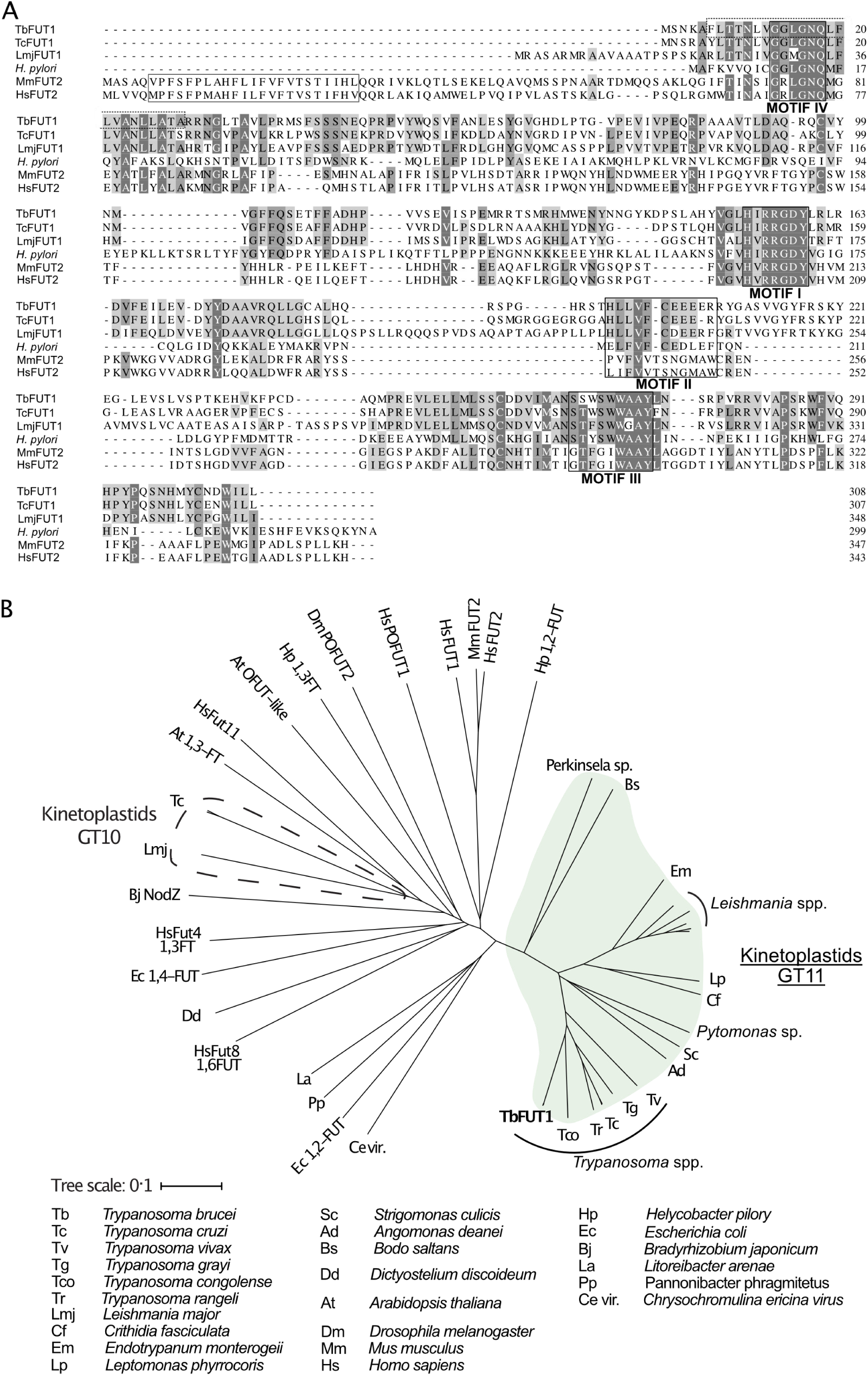
Amino acid sequence and phylogenetic analyses of TbFUT1. *A*. Sequence alignment of TbFUT1 and other GT11 family FUTs shows that TbFUT1 (Tb927.9.3600) lacks a conventional N-terminal type-2 membrane protein transmembrane domain (TM, grey box) but contains conserved motifs I-IV (*black boxes*). A putative TbFUT1 TM (*dashed box*) overlaps with motif IV but does not align with the eukaryotic FUT TM and is most likely part of a cleavable *N*-terminal mitochondrial targeting sequence (residues 1-52). Sequences used in the alignment: *T. cruzi* (TcCLB.506893.90), *L. major* (LmjF01.0100), *H. pylori* (AAC99764), *H. sapiens* FUT2 (AAC24453) and *M. musculus* FUT2 (AAF45146). *B*. A selection of known and predicted fucosyltransferase protein sequences were aligned using ClustalΩ (Sievers et al., 2011) and the unrooted phylogram shown was generated by iTOL (itol.embl.de) (Letunic and Bork, 2016). Single homologues of the GT11 TbFUT1 were found in all the kinetoplastids (marked in *green*) and, collectively, these sequences form a clade distant from other fucosyltransferase sequences, including the other kinetoplastid GT10 fucosyltransferases (marked by a *dashed line*). No TbFUT1 homologues were found in apicomplexan parasites or in Euglena.

### Recombinant expression of TbFUT1

The *TbFUT1* ORF was amplified from *T. brucei* 427 genomic DNA and cloned in the pGEX6P1 expression vector. The resulting construct (pGEX6P1-GST-PP-*TbFUT1*) encoded for the *TbFUT1* ORF with a glutathione-S-transferase (GST) tag at its N-terminus and a PreScission™ Protease (PP) cleavage site between the two protein-encoding sequences. Sequencing confirmed what was subsequently deposited at TriTrypDB (Tb427_090021700) and identified two amino acid differences between FbFUT1 in the 927 and 427 strains (L185V and T232A).

The pGEX6P1-GST-PP-*TbFUT1* construct was expressed in *E. coli* and the fusion protein purified as described in Material and Methods. The identities of the two higher molecular weight bands (Fig. S1, lane 8) were determined by peptide mass fingerprinting. The most abundant band was identified as TbFUT1, while the fainter band was identified as a subunit of the *E. coli* GroEL chaperonin complex. The apparent molecular weight of GST-PP-TbFUT1 chimeric protein (57 kDa) was consistent with the predicted theoretical molecular weight (58.1 kDa).

### Recombinant TbFUT1 is active in vitro

The activity of recombinantly expressed GST-TbFUT1 fusion protein was tested by incubation with GDP-[^3^H]Fuc, as a donor, and a panel of commercially available mono-to octasaccharides (Table 1) selected from the literature as possible α-1,2-FUT substrates (Li et al., 2008; Wang et al., 1999; Zhang et al., 2010). The effectiveness of each acceptor was evaluated based on the presence/absence and intensities of the TLC bands corresponding to the radiolabelled reaction products (Fig. 2 and Table 1). GST-TbFUT1 showed best activity with Galβ1,3GlcNAc (LNB) (Fig. 2, lane 2 and 5) and its β-O-Methyl glycoside (Fig. 2, lane 23). Other larger oligosaccharides containing Galβ1,3GlcNAcβ1-O-R as a terminal motif (LNT and LNH) were also good acceptors (Fig. 2, lanes 13 and 16), with the exception of iLNO (Fig. 2, lane 15). Lactose was also recognized (Fig. 2, lane 1), while LacNAc and the LacNAc-terminating branched hexasaccharide LNnH were weak acceptors (Fig. 2, lanes 3 and 12). Interestingly, TbFUT1 was also able to transfer fucose to 3’-fucosyllactose, albeit inefficiently (Fig. 2, lane 18), whereas no transfer could be seen to Galβ1,6GlcNAc (Fig. 2, lane 17) or to free Gal or β-Gal-O-methyl (Fig. 2, lanes 11 and 22). As expected, no products were observed when acceptor oligosaccharides were omitted from the reaction (Fig. 2, lanes 4 and 6). To confirm the detected activities were specific to the recombinant GST-TbFUT1, and not due to some co-purifying endogenous *E. coli* contaminant, the assay was also performed using material prepared from *E. coli* expressing the empty pGEX6P1 vector. No transfer of radiolabelled fucose could be observed under these conditions (Fig. 2, lanes 7-9).

**FIGURE 2.**
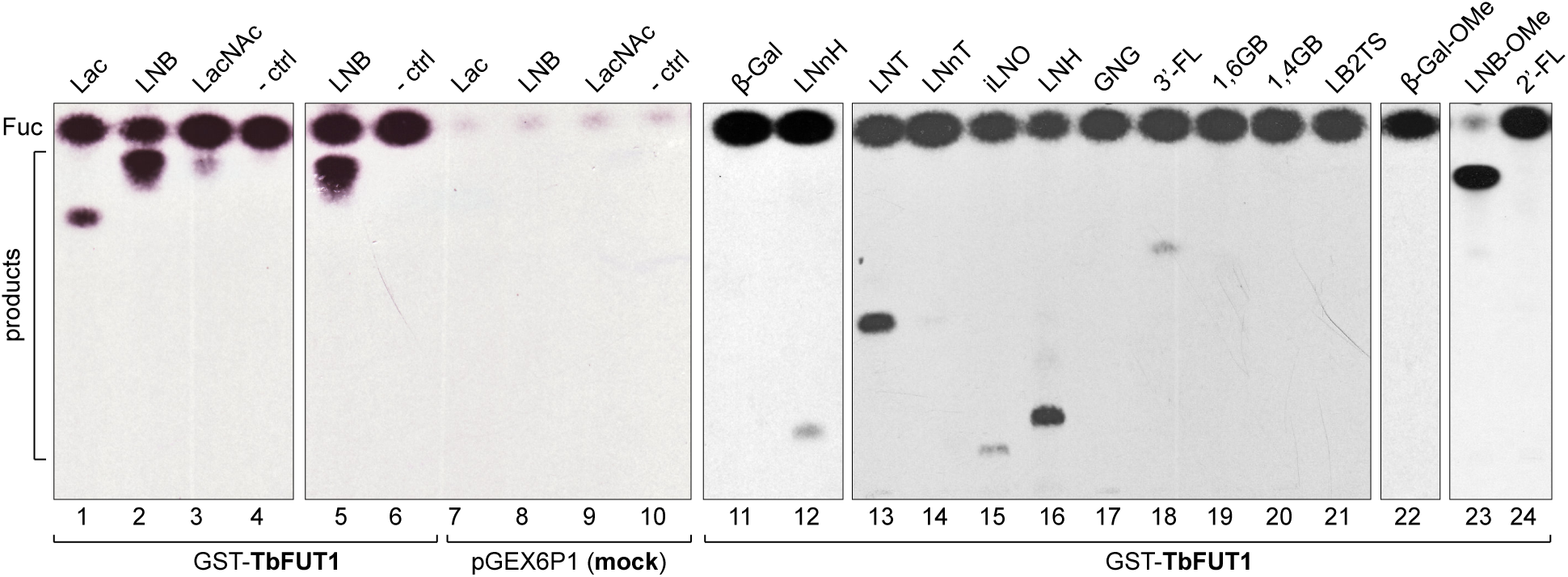
Recombinant GST-TbFUT1 transfers [^3^H]Fuc to a variety of sugar acceptors. Each assay used 2 μg of purified GST-TbFUT1, GDP-[^3^H]Fuc and 1 mM of acceptor. Reaction products were desalted and separated by silica HPTLC, and detected by fluorography. LNB, LNB-OMe, and LNB-terminating structures were the best acceptors tested. The acceptor abbreviations above each lane are defined in Table 1. *- ctrl*: negative control reaction without acceptor (Lanes 4 and 6) or with buffer alone (lane 10).

**Table 1.**
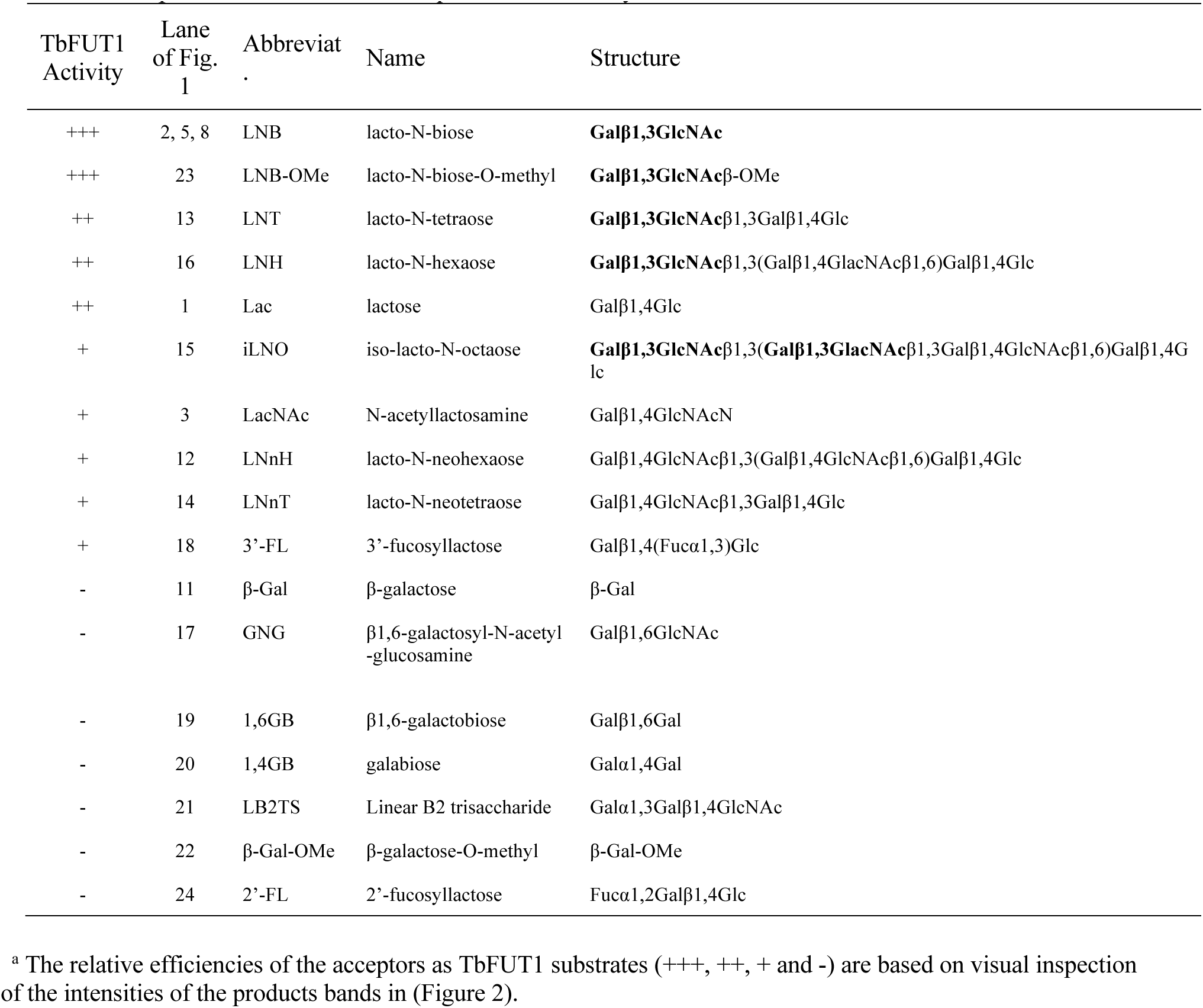
Acceptor substrates and semi-quantitative fucosyltransferase activities.

A band with the same mobility as free fucose was observed in all assay reactions and was considerably stronger in the presence of the GST-TbFUT1 preparation (Fig. 2, lanes 1-6 and 11-24) than when GDP-[^3^H]Fuc was incubated with reaction buffer alone (Fig. 2, lane 10) or in the presence of the material purified from the *E. coli* cells transformed with the empty vector (Fig. 2, lanes 7-9). These data suggest that TbFUT1 has a significant propensity to transfer Fuc to water. Interestingly, one of the substrates (LNB-O-Me; Galβ1,3GlcNAcβ1-O-methyl) suppressed the amount of free Fuc produced in the reaction (Fig. 2, lane 23), suggesting that this glycan may bind more tightly to the TbFUT1 acceptor site than the other oligosaccharides tested and thus prevent the transfer of Fuc from GDP-Fuc to water. *In vitro* high sugar nucleotide hydrolysis activity has been previously described for at least one member of the GT11 family (Zhang et al, 2010).

Inverting α-1,2 and α-1,6-FUTs do not usually require divalent cations for their activity (Beyer and Hill, 1980; Kamińska et al., 2003; Li et al., 2008; Pettit et al., 2010). To study the divalent cation dependence of TbFUT1, the assay was repeated in buffer without divalent cations or containing EDTA. No change in activity was observed in either case, indicating TbFUT1 does not require divalent cations for its activity (Fig. S2).

### Characterization of the TbFUT1 reaction product

The glycan reaction products were structurally characterized to determine the anomeric and stereochemical specificity of TbFUT1. Initially, we performed exoglycosidase and/or acid treatment of the radiolabelled reaction products (recovered by preparative TLC) utilizing Lac, LacNAc and LNB as substrates. The tritium label ran with the same mobility as authentic Fuc after acid hydrolysis of all three products (Fig. S3*A* and *C*) and after *Xanthomonas manihotis* α-1,2-fucosidase digestion of the Lac and LNB products (Fig. S3*B* and *C*). These data suggest that [^3^H]Fuc was transferred in α1,2 linkage to the acceptor disaccharides.

To obtain direct evidence, we performed a large-scale activity assay using LNB-O-Me as an acceptor and purified the reaction product by normal phase HPLC. Fractions containing the putative fucosylated trisaccharide product (Fig. S4) were pooled and analysed for their neutral monosaccharide content, which showed the presence of Fuc, Gal and GlcNAc. The purified reaction products were permethylated and analysed by ESI-MS (Fig. 3*A*), which confirmed that the main product was a trisaccharide of composition dHex_1_Hex_1_HexNAc_1_. The MS/MS spectrum was also consistent with the dHex residue being attached to the Hex, rather than HexNAc, residue (Fig. 3*B*). Subsequently, partially methylated alditol acetates (PMAAs) were generated from the purified trisaccharide product and analysed by GC-MS. This analysis identified derivatives consistent with the presence of non-reducing terminal-Fuc, 2-*O*-substituted Gal and 3-*O*-substituted GlcNAc (Fig. S5 and Table 2), consistent with Fuc being linked to position 2 of Gal. The GC-MS methylation linkage analysis also revealed a trace of 2-*O*-substituted Fuc in the sample which, together with the observation that 3’-FL can act as a weak substrate (Fig. 2, lane 18 and Table 1), may suggest that TbFUT1 can also form Fucα1,2Fuc linkages.

**FIGURE 3.**
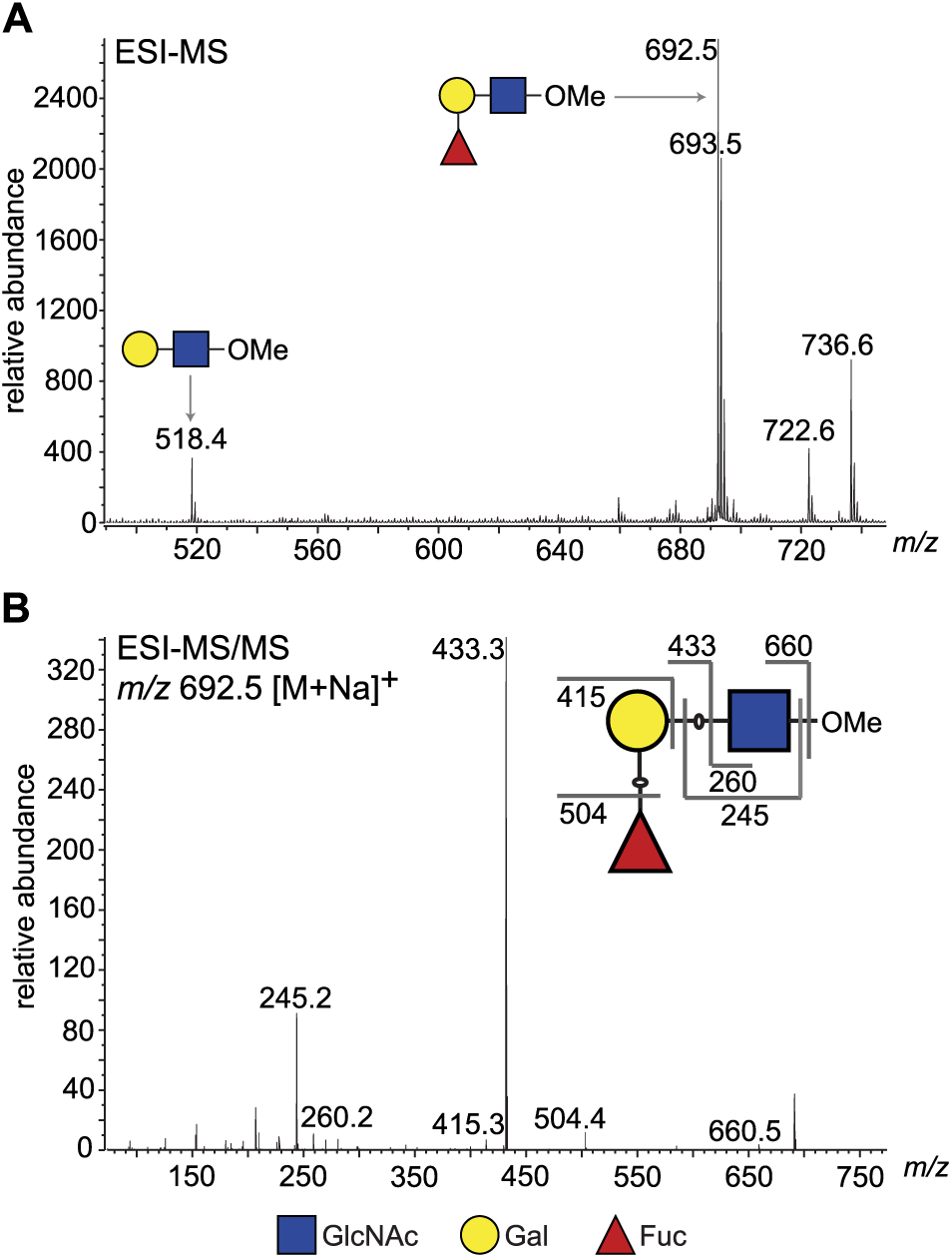
ESI-MS and ESI-MS/MS of TbFUT1 *in vitro* reaction product. *A*. ESI-MS of the purified and permethylated reaction product. The ion at *m/z* 692.5 is consistent with the [M + Na]^+^ ion of a permethylated trisaccharide of composition dHex_1_Hex_1_HexNAc_1_. Some of the unmodified acceptor (Hex_1_HexNAc_1_) was also observed (*m/z* 518.4). *B*. MS/MS product ion spectrum of *m/z* 692.5. The collision-induced fragmentation pattern indicated that the dHex (Fuc) residue was linked to the Hex (Gal) and not to the HexNAc (GlcNAc) residue.

**Table 2.**
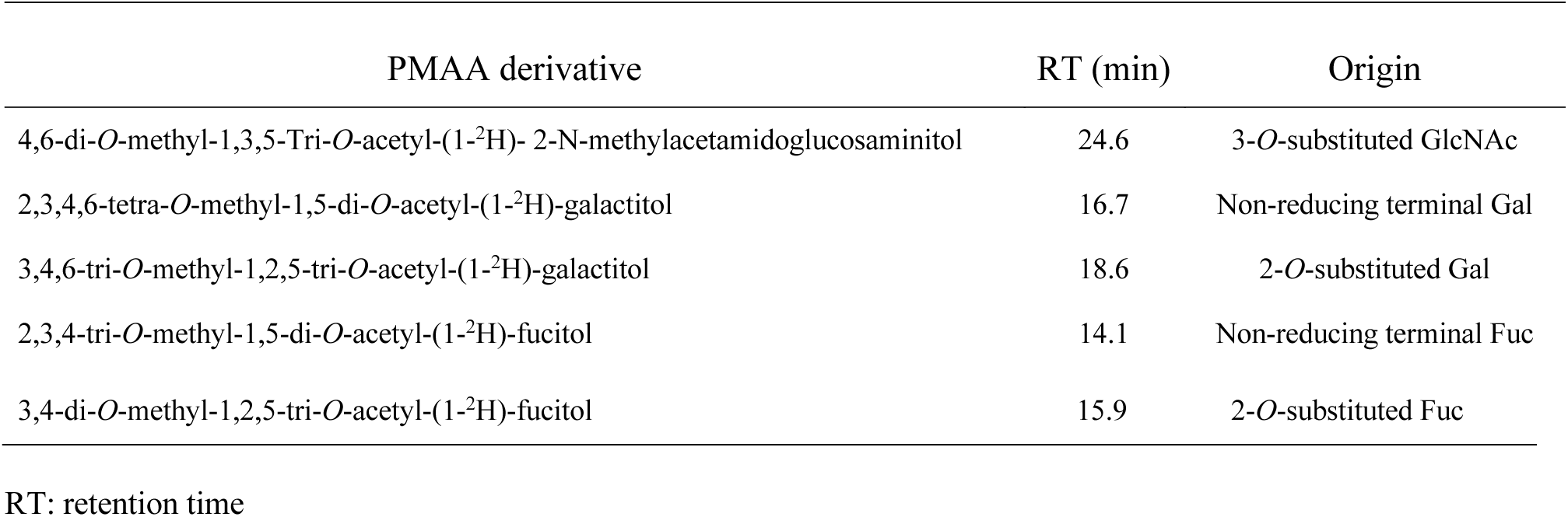
PMAAs derivatives identified by GC-MS methylation linkage analysis of the purified TbFUT1 reaction product.

| PMAA derivative | RT (min) | Origin |
| --- | --- | --- |
| 4,6-di-*O*-methyl-1,3,5-Tri-*O*-acetyl-(1-$^2\text{H}$)- 2-N-methylacetamidoglucosaminitol | 24.6 | 3-*O*-substituted GlcNAc |
| 2,3,4,6-tetra-*O*-methyl-1,5-di-*O*-acetyl-(1-$^2\text{H}$)-galactitol | 16.7 | Non-reducing terminal Gal |
| 3,4,6-tri-*O*-methyl-1,2,5-tri-*O*-acetyl-(1-$^2\text{H}$)-galactitol | 18.6 | 2-*O*-substituted Gal |
| 2,3,4-tri-*O*-methyl-1,5-di-*O*-acetyl-(1-$^2\text{H}$)-fucitol | 14.1 | Non-reducing terminal Fuc |
| 3,4-di-*O*-methyl-1,2,5-tri-*O*-acetyl-(1-$^2\text{H}$)-fucitol | 15.9 | 2-*O*-substituted Fuc |

RT: retention time

The purified TbFUT1 reaction product was also exchanged into deuterated water (^2^H_2_O) and analysed by one-dimensional ^1^H-NMR and two-dimensional ^1^H-ROESY (Rotating frame Overhouser Effect SpectroscopY). The ^1^H-NMR spectrum showed a doublet at about 5.1 ppm, consistent with the signal from the proton on the anomeric carbon (H_1_) of an α-Fuc residue (Fig. 4*A*).

**FIGURE 4.**
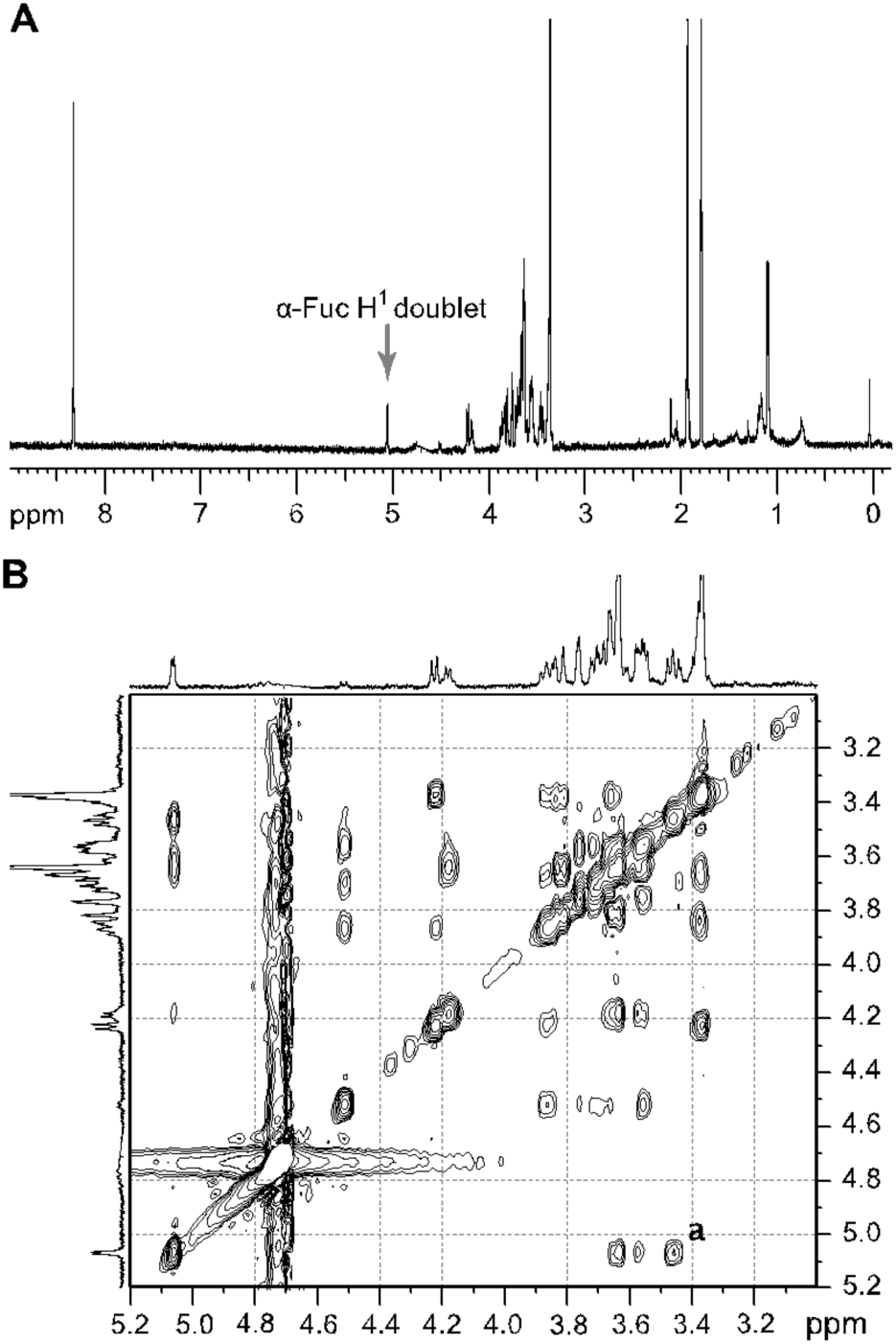
^1^H NMR and ^1^H ROESY spectra of the TbFUT1 reaction product. *A*. One-dimensional ^1^H-NMR spectrum. The *arrow* points to the α-Fuc H_1_ doublet. *B*. Enlargement of the 3.2-5.1 ppm region of the two-dimensional ^1^H ROESY spectrum. ***a*** indicates the crosspeak resulting from a through-space connectivity between α-Fuc H_1_ and β-Gal H_2_.

A characteristic doublet for the anomeric proton of a β-Gal residue was also observed at 4.5 ppm. In the ^1^H-ROESY spectrum, a cross-peak (labelled *a*) could be observed indicating a through-space connectivity between the H_1_ of α-Fuc and the H_2_ of a β-Gal, consistent with a Fucα1,2Gal linkage in the TbFUT1 reaction product (Fig. 4*B*). The chemical shifts that could be clearly assigned by either one-dimensional ^1^H-NMR or two-dimensional ^1^H-ROESY are listed in Table 3.

**Table 3.** ^1^H and ^1^H ROESY chemical shift assignments for the purified TbFUT1 reaction product

| Residue | $\text{H}_1$ | $\text{H}_2$ | $\text{H}_3$ | $\text{H}_4$ | $\text{H}_5$ | $\text{H}_{6/6'}$ | NAc |
| --- | --- | --- | --- | --- | --- | --- | --- |
| αFuc | 5.05 (J=4 Hz) | 3.57 | 3.67 | 3.63 | 4.2 | 1.1 | |
| βGal | 4.5 | 3.45 | 3.55 | 3.89 | ND | ND | |
| βGlcNAc | ND | 3.63 | ND | 3.4 | ND | 3.78/3.89 | 2.1 |

J: coupling constant
ND: chemical shift could not be clearly assigned.

Taken together, these data unambiguously define the structure of the TbFUT1 reaction product with GDP-Fuc and LNB-O-Me as Fucα1,2Galβ1,3GlcNAcβ1-O-CH_3_ which, in turn, defines TbFUT1 as having a GDP-Fuc : βGal α-1,2 fucosyltransferase activity with an apparent preference for a Galβ1,3GlcNAcβ1-O-R acceptor motif.

### Generation of TbFUT1 conditional null mutants in procyclic and bloodstream form T. brucei

Semi-quantitative RT-PCR showed that *TbFUT1* mRNA was present in both bloodstream form and procyclic form *T. brucei* (data not shown). We therefore sought to explore TbFUT1 function in both lifecycle stages by creating *TbFUT1* conditional null mutants. The strategies used to generate the mutants are described in (Fig. 4). The creation of these mutants was possible because genome assembly indicated *TbFUT1* to be present as a single copy per haploid genome, and Southern blot analysis using a *TbFUT1* probe was consistent with this prediction (Fig. S6). In procyclic cells (Fig. 5, left panel), the first *TbFUT1* allele was replaced by homologous recombination with linear DNA containing the puromycin resistance gene (*PAC*) flanked by about 500 bp of the *TbFUT1* 5’- and 3’-UTRs. After selection with puromycin, an ectopic copy of *TbFUT1,* under the control of a tetracycline-inducible promoter, was introduced in the ribosomal DNA (rDNA) locus using phleomycin selection. Following induction with tetracycline, the second allele was replaced with the *BSD* gene by homologous recombination, generating the final procyclic form *ΔTbFUT1::PAC/TbFUT1^Ti^/ΔTbFUT1::BSD* conditional null mutant cell line (PCF *TbFUT1* cKO). In bloodstream form cells (Fig. 5, middle panel), an ectopic copy of *TbFUT1* carrying a C-terminal MYC_3_ epitope tag under the control of a tetracycline-inducible promoter was first introduced into the ribosomal DNA (rDNA) locus using phleomycin selection. Following cloning and induction with tetracycline, the first *TbFUT1* allele was then targeted for homologous recombination with linear DNA containing the hygromycin resistance gene (*HYG*) flanked by about 1200 bp of the *TbFUT1* 5’- and 3’-UTRs. After selection with hygromycin, Southern blotting revealed that gene conversion had taken place and that both *TbFUT1* alleles had been replaced by *HYG* yielding a bloodstream form *TbFUT1-MYC_3_^Ti^/ΔTbFUT1::HYG*/*ΔTbFUT1::HYG* conditional null mutant cell line (BSF *TbFUT1*-MYC_3_ cKO). Southern blotting data confirming the genotypes of these mutants are shown in (Fig. S6). The BSF cell line was also used to generate a *TbFUT1^Ti^/ΔTbFUT1::HYG*/*ΔTbFUT1::HYG* conditional null mutant cell line by *in situ* homologous recombination of the tetracycline inducible *TbFUT1*-MYC_3_ copy, converting it to an untagged *TbFUT1* gene and generating BSF *TbFUT1* cKO (Fig. 5, right panel).

**FIGURE 5.**
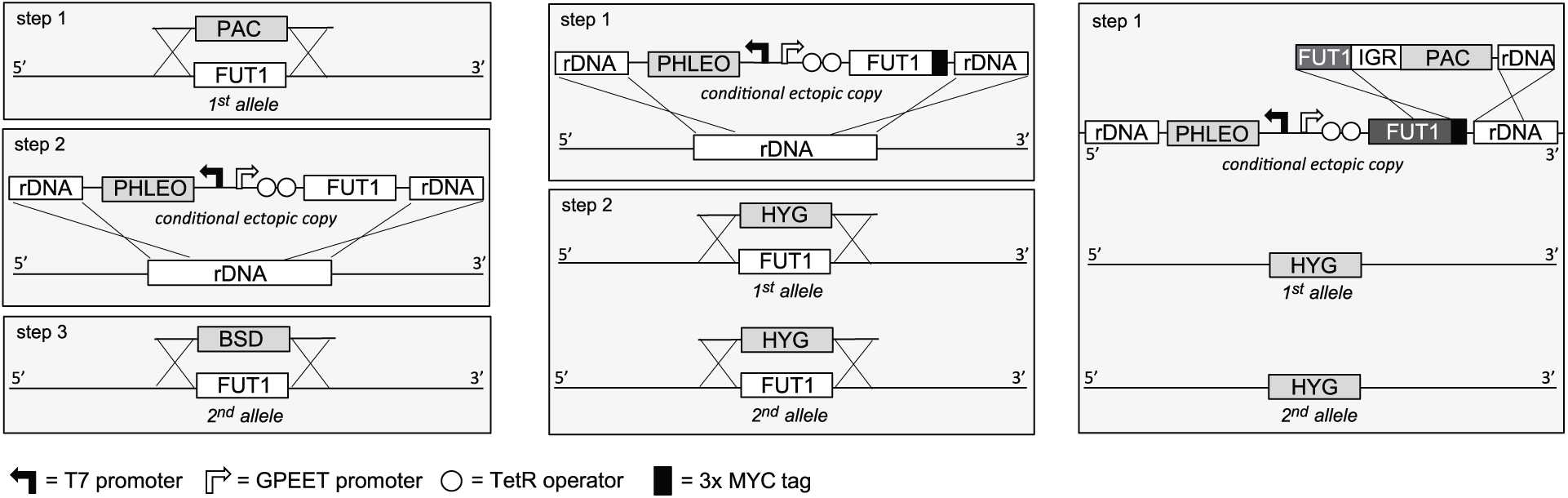
Cloning strategies for the creation of the *TbFUT1* conditional null mutants. *Left panel*: To create the procyclic form conditional null mutant (PCF *TbFUT1* cKO) the first *TbFUT1* allele was replaced by *PAC*, an ectopic tetracycline-inducible copy of the *TbFUT1* gene was introduced into the ribosomal DNA locus and the second *TbFUT1* allele was replaced by *BSD*. *Middle panel*: To create the bloodstream form conditional null mutant (BSF *TbFUT1-MYC_3_* cKO) an ectopic tetracycline-inducible copy of the *TbFUT1* gene with a MYC_3_ tag was first introduced into the ribosomal DNA locus. Both *TbFUT1* alleles were subsequently replaced by *HYG* through homologous recombination followed by gene conversion. *Right panel:* To create the untagged bloodstream form cKO (BSF *TbFUT1* cKO), the BSF *TbFUT1-MYC_3_* cKO mutant (*middle panel*) was modified by homologous recombination with a construct that removed the C-terminal MYC_3_ tag under *PAC* selection.

### TbFUT1 is essential to procyclic and bloodstream form T. brucei

Procyclic and bloodstream form *TbFUT1* conditional null mutants were grown under permissive (plus tetracycline) or non-permissive (minus tetracycline) conditions. The PCF *TbFUT1* cKO cells grown under non-permissive conditions showed a clear reduction in the rate of cell growth after 6 days, eventually dying after 15 days (Fig. 6*A*).

**FIGURE 6.**
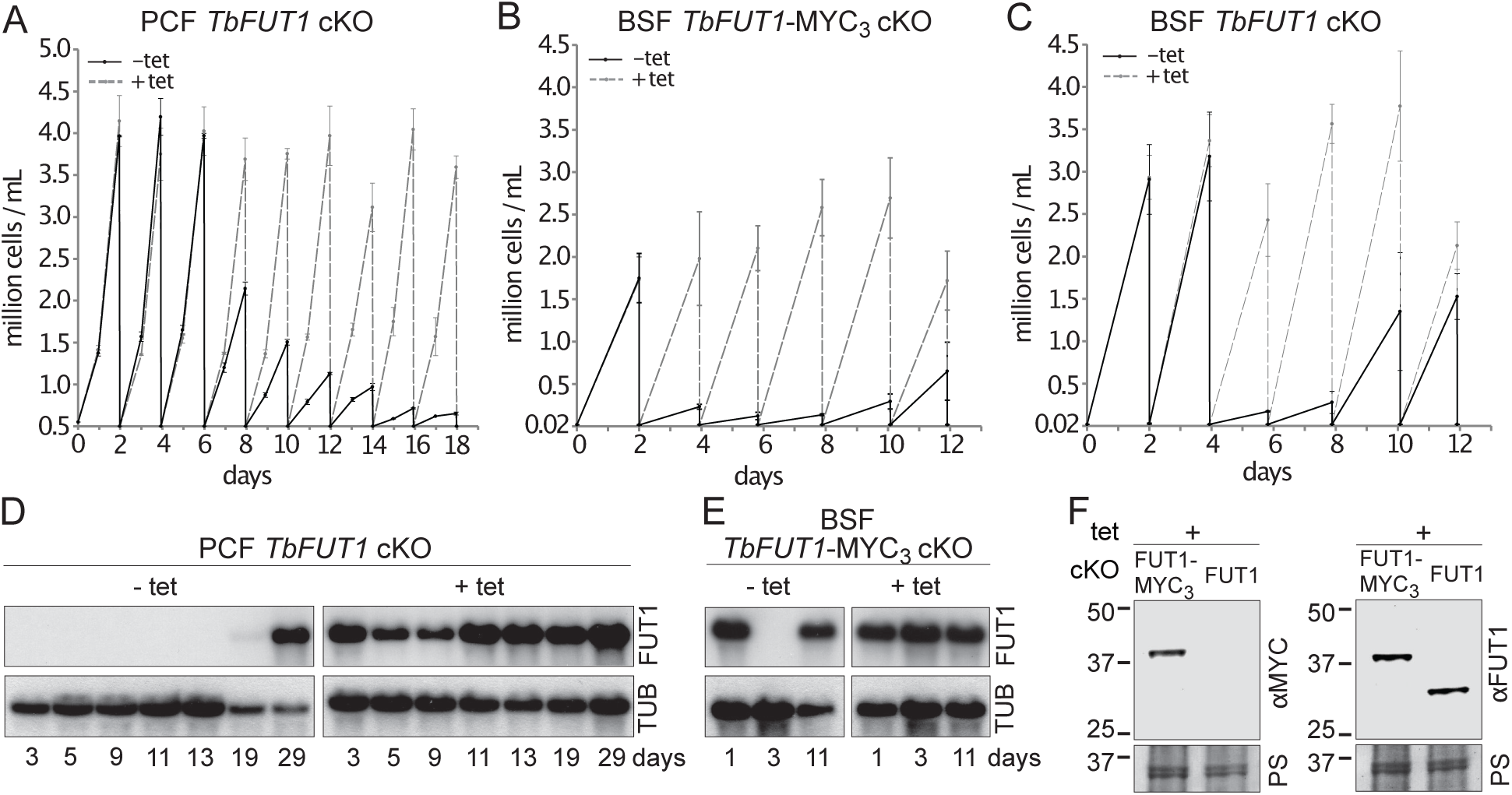
TbFUT1 is essential for procyclic and bloodstream form cell growth *in vitro.* The cell numbers (± standard deviation) for TbFUT1 cKO under permissive (plus tetracycline, *dotted line*) and non-permissive (minus tetracycline, *solid line*) conditions are shown for three procyclic (*A*) and bloodstream form (*C*) clones, as well as for three bloodstream clones carrying a tetracycline-inducible ectopic *TbFUT1* gene with a C-terminal MYC_3_ tag (*B*). *D-E*. Corresponding *TbFUT1* mRNA levels were determined by Northern blots. Alpha-Tubulin (TUB) was used as a loading control. *F*. TbFUT1-MYC_3_ and untagged TbFUT1 are detected by Western blot analysis in the respective bloodstream form cKO cell lines under permissive conditions (+ Tet). The *left panel* shows an anti-MYC (αMYC) blot and the *right panel* an anti-recombinant TbFUT1 antibody (αFUT1) blot. Membranes were stained with Ponceau S (PS) to ensure equal loading.

The BSF *TbFUT1* cKO cells grew like wild type cells under permissive conditions, whether or not the expressed TbFUT1 had a C-terminal MYC_3_ tag, and under non-permissive conditions also showed a clear reduction in the rate of cell growth after 2-4 days, dying after 3-5 days (Fig. 6*B* and *C*). These growth phenotypes are very similar to those described for procyclic and bloodstream form *TbGMD* conditional null mutants that cannot synthesise GDP-Fuc under non-permissive conditions (Fig. S7) (Turnock et al., 2007). This is consistent with the hypothesis that TbFUT1 may be the only enzyme that utilizes GDP-Fuc, or at least that it is the only FUT transferring fucose to essential acceptors. Further evidence that TbFUT1 is essential for procyclic and bloodstream form growth was obtained from Northern blots (Fig. 6*D* and *E*). These show that *TbFUT1* mRNA levels are undetectable for several days after the removal of tetracycline, but that growth resumes only when some cells escape tetracycline control after about 29 days (procyclic form) and 11 days (bloodstream form). Escape from tetracycline control after several days is typical of conditional null mutants for essential trypanosome genes (Roper et al., 2002). Evidence for the expression of the MYC_3_ tagged TbFUT1 protein in the BSF *TbFUT1*-MYC_3_ cKO cell line and of unmodified TbFUT1 in the BSF *TbFUT1* cKO cell line under permissive conditions is shown in (Fig. 6*F*).

From a morphological point of view, both procyclic form *TbGMD* and *TbFUT1* conditional null mutants grown under non-permissive conditions showed an increase in average cell volume, due to increased cell length, concomitant with the start of the cell growth phenotype (Fig. S8*A*). However, we were unable to reproduce the flagellar detachment phenotype previously reported for the PCF *TbGMD* cKO grown in non-permissive conditions (Turnock et al., 2007), nor was such a phenotype observed in the PCF *TbFUT1* cKO parasites, either by scanning electron microscopy or immunofluorescence (Fig. S8*C* and S9). The percentage of cells displaying flagellar detachment (1.5-2 %) in both null mutants grown in non-permissive conditions (Fig. S8*B*) was consistent with what has previously been reported for wild type cells (LaCount et al., 2002). Additionally, we could observe no defect in cell motility in either PCF *TbGMD* or PCF *TbFUT1* cKO grown in non-permissive conditions (Fig. S10).

### TbFUT1 localizes to the parasite mitochondrion

The BSF *TbFUT1*-MYC_3_ cKO cell line, grown under permissive conditions, was stained with anti-MYC antibodies and produced a pattern suggestive of mitochondrial localization. This was confirmed by co-localization with MitoTracker^TM^ (Fig. 7*A*). However, when TbFUT1 was introduced into wild type cells fused with an HA_3_ epitope tag at the N-terminus, either with or without a C-terminal MYC_3_-tag (constructs pLEW100:HA_3_-FUT1-MYC_3_ and pLEW100:HA_3_-FUT1), the tagged protein co-localized with GRASP, a marker of the Golgi apparatus (Fig. 7*B* and *C*). In these cases, we suspect that N-terminal tagging has disrupted mitochondrial targeting, by obscuring the N-terminal mitochondrial targeting sequence. Indeed, no mitochondrial targeting motif was predicted *in silico* for N-terminal HA_3_ tagged TbFUT1. Nevertheless, since the mitochondrial localization of a fucosyltransferase is unprecedented, we elected to raise polyclonal antibodies against recombinant TbFUT1 to further assess its subcellular location. To do so, an N-terminally His_6_-tagged Δ_32_TbFUT1 protein was expressed, re-solubilized from inclusion bodies and used for rabbit immunization. The IgG fraction was isolated on immobilized protein-A and the anti-TbFUT1 IgG sub-fraction affinity purified on immobilized recombinant GST-TbFUT1 fusion protein. To further ensure mono-specificity of the antibodies to TbFUT1, the resulting fraction was adsorbed against a concentrated cell lysate of the PCF *TbFUT1* cKO mutant grown for 9 days under non-permissive conditions. The resulting highly-specific polyclonal antibody was used to detect TbFUT1 expression in wild type bloodstream form cells as well as in BSF and PCF *TbFUT1* cKO cells under permissive and non-permissive conditions (Fig. 8*A-C*). Anti-TbFUT1 antibodies co-localized with MitoTracker^TM^ staining in the wild type cells and in the conditional null mutants under permissive conditions. No signal for the anti-TbFUT1 antibodies was seen under non-permissive conditions, confirming the specificity of the polyclonal antibody. Taking the possibility of a dual Golgi/mitochondrial localization into account, TbFUT1 localization was also assessed in bloodstream form cells ectopically expressing TbGnTI-HA_3_ as an authentic Golgi marker (Damerow et al., 2014). No co-localization between TbGnTI-HA_3_ and anti-TbFUT1 was observed, suggesting that TbFUT1 is either exclusively or predominantly expressed in the parasite mitochondrion (Fig. 8*D*).

**FIGURE 7.**
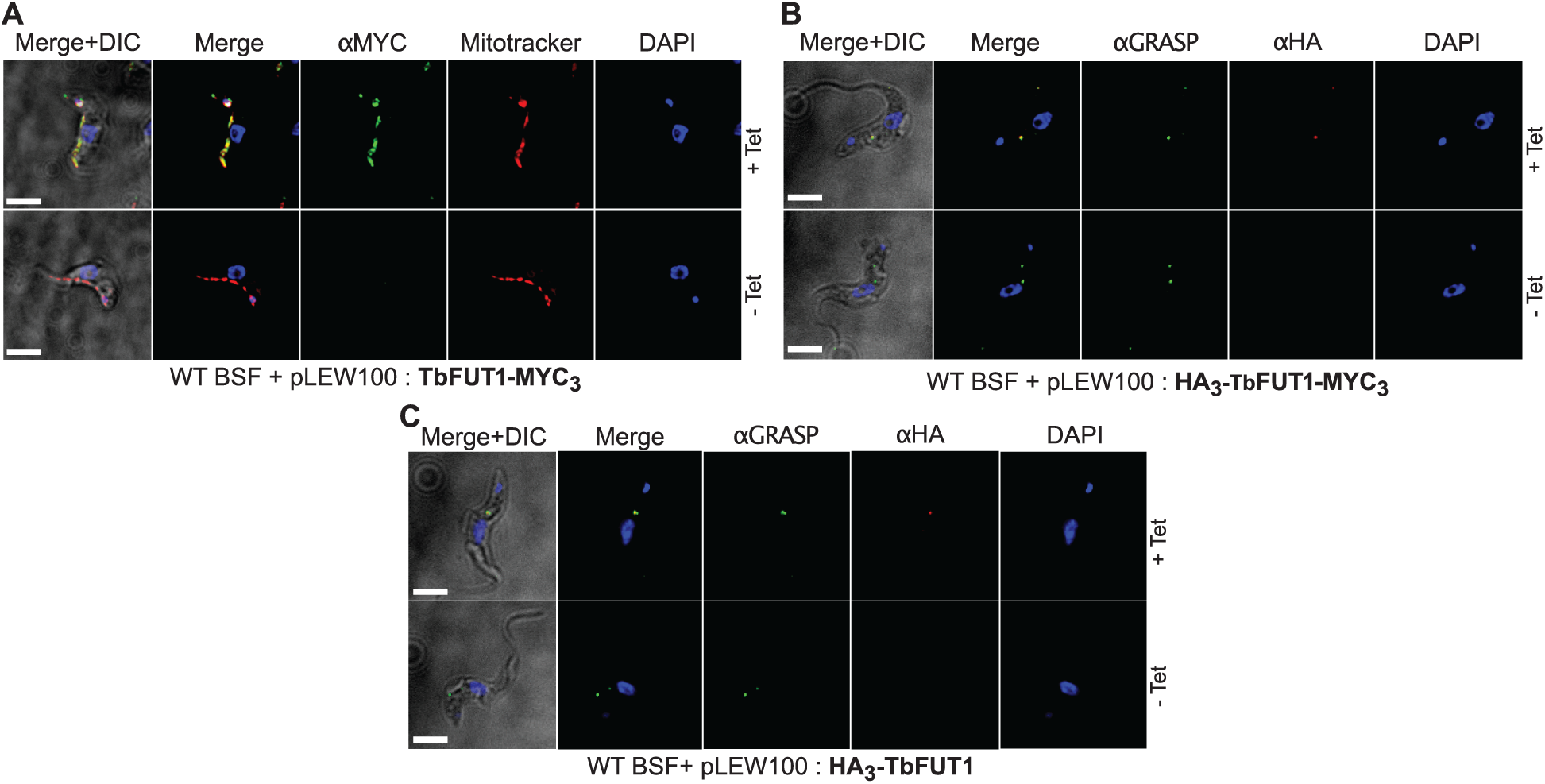
C- and N-terminal tagging of TbFUT1 result in mitochondrial and Golgi apparatus localization, respectively. *A.* Bloodstream form (BSF) cKO parasites expressing tet-inducible C-terminally tagged TbFUT1-MYC_3_ were imaged under permissive (+Tet) and non-permissive (-Tet) conditions by DIC and fluorescence microscopy after staining with anti-MYC, MitoTracker^TM^, and DAPI. Comparable patterns were observed for anti-MYC and MitoTracker^TM^, suggesting TbFUT1-MYC_3_ localizes to the mitochondrion. *B-C*. IFA of BSF cKO parasites expressing a tet-inducible N-terminally tagged HA_3_-TbFUT1-MYC_3_ (*B*) or HA_3_-TbFUT1 (*C*) after labelling with anti-HA, anti-GRASP, and DAPI suggests a Golgi apparatus location for both HA_3_-TbFUT1-MYC_3_ and HA_3_-TbFUT1. The absence of anti-MYC (*A*) or anti-HA (*B-C*) staining under non-permissive conditions confirms the specificity of the labelling for the respective TbFUT1 fusion proteins. Scale bars: 3 μm.

**FIGURE 8.**
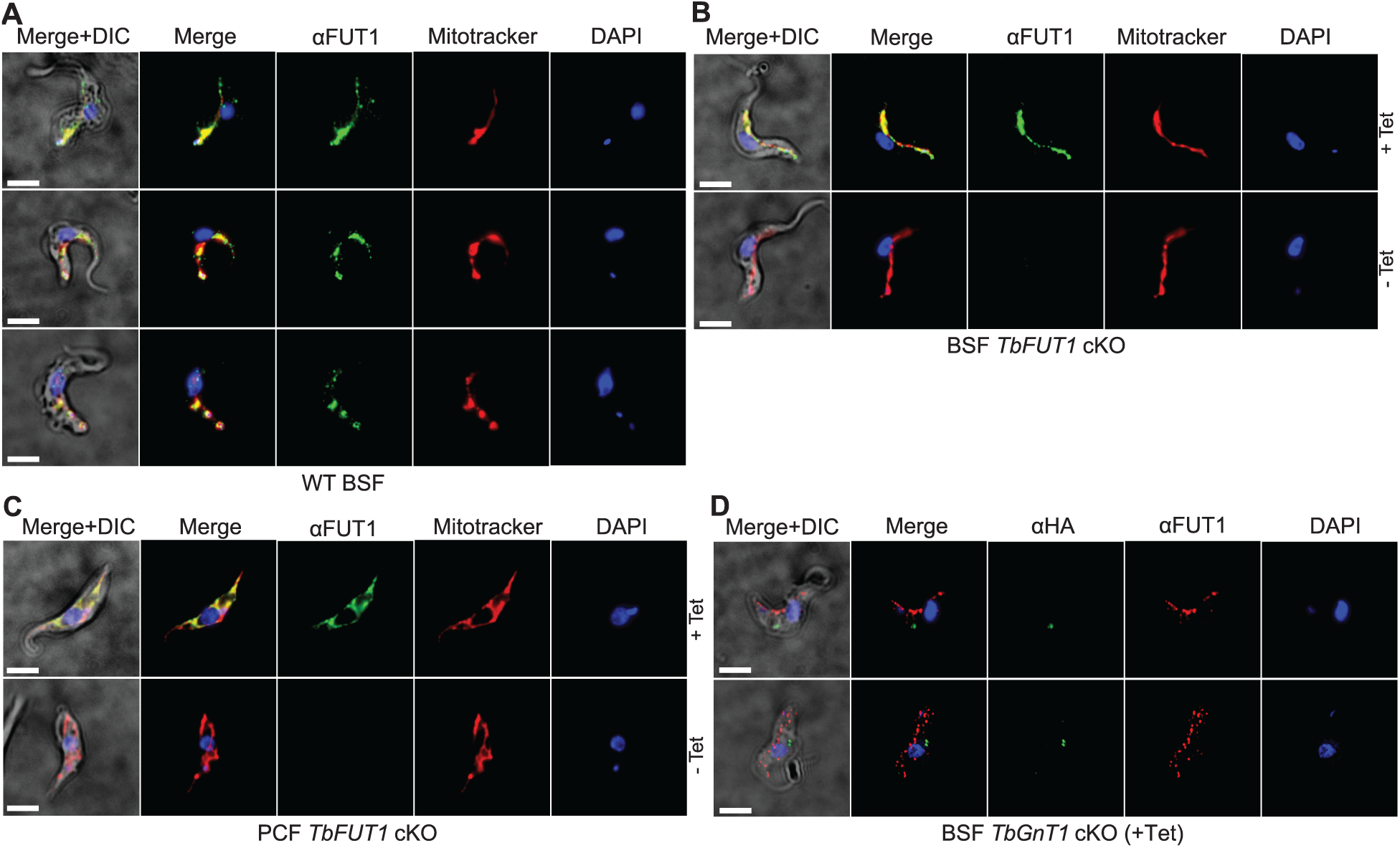
Antibodies to the recombinant protein localize TbFUT1 to the mitochondrion. *A*. IFA of wild type bloodstream form (BSF) trypanosomes after staining with affinity purified anti-TbFUT1 (αFUT1) MitoTracker^TM^ and DAPI. Comparable patterns were observed for αFUT1 and MitoTracker^TM^, suggesting TbFUT1 localizes to the mitochondrion. *B-C*. Bloodstream (*B*) and procyclic (*C*) form *TbFUT1* conditional null mutants were imaged under permissive (+Tet) and non-permissive (-Tet) conditions. In both cases the tetracycline-inducible TbFUT1 pattern is consistent with a mitochondrial localization. *D.* BSF trypanosomes induced to express a C-terminally tagged known Golgi glycosyltransferase (TbGnTI-HA_3_) were imaged after staining with αFUT1, anti-HA and DAPI, as indicated. The merged images of two representative cells suggest no significant co-localization between native TbFUT1 and the Golgi-localized TbGnT1. Scale bars: 3 μm.

### Deletion of TbFUT1 disturbs mitochondrial activity

Bloodstream form wild type and BSF *TbFUT1* cKO cells, grown with and without tetracycline for 5 days, were stained with antibodies to mitochondrial ATPase and with MitoTracker^TM^. As expected ATPase and MitoTracker^TM^ co-localized in wild type cells and in the mutant under permissive conditions (Fig. 9, top panels). However, under non-permissive conditions the few remaining viable cells showed significantly diminished MitoTracker^TM^ staining, indicative of reduced mitochondrial membrane potential, and a reduction in ATPase staining, suggesting that TbFUT1 is in some way required for mitochondrial function (Fig. 9 lower panels, and Fig. S11).

**FIGURE 9.**
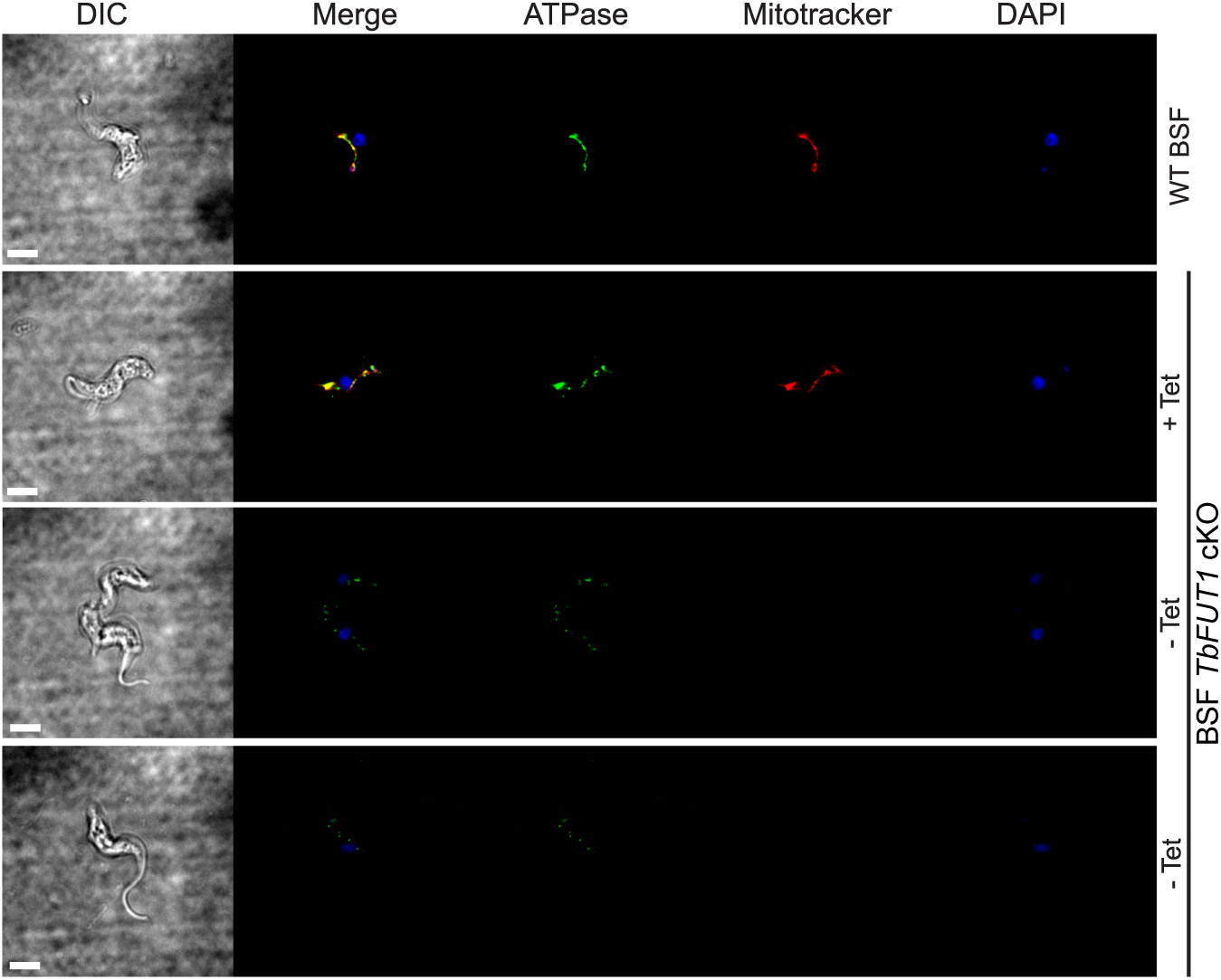
Absence of TbFUT1 disturbs mitochondrial activity. Bloodstream form (BSF) wild type and *TbFUT1* cKO parasites were cultured for 5 days under permissive (+ Tet) and non-permissive (- Tet) conditions, fixed and labelled with Mitotracker^TM^ for mitochondrial potential and with anti-mitochondrial ATPase antibody. In mutants grown in non-permissive conditions (*lower panels*) both ATPase and Mitotracker staining are strongly reduced, suggesting reduced mitochondrial functionality. Scale bar: 3 μm.

### TbFUT1 can replace the homologous essential mitochondrial fucosyltransferase in Leishmania major

The essentiality and mitochondrial location of the orthologous protein in *L. major*, LmjFUT1, was reported in (Guo et al. 2021). Here we asked whether TbFUT1 could functionally substitute for LmjFUT1, using a plasmid shuffle approach (Guo et al., 2017; Murta et al., 2009). We started with an *L. major* homozygous *fut1*-null mutant expressing *LmjFUT1*from the episomal pXNGPHLEO vector which additionally expresses GFP (Guo et al., 2017; Murta et al., 2009) and transfected it with an episomal construct pIR1NEO-*TbFUT1* (±HA) expressing WT or HA-tagged *TbFUT1*. Following transfection, parasites were grown briefly in the absence of any selective drug, and single cells were sorted for GFP expression and selected for phleomycin resistance, or GFP loss accompanied by G418 (*NEO*) resistance (Fig. 10*A*). In these studies, 85% (82/96) of the GFP-positive (GFP+) clonal lines (positive control) grew when placed in microtiter wells, while 81% (156/192) of the GFP-negative (GFP-) sorted cells likewise grew (Fig. 10*A*). Control PCR studies confirmed the loss of *TbFUT1* in the GFP-clones (not shown). The similar survival of cells bearing *Lmj* or *TbFUT1* provides strong evidence that the TbFUT1 can fully satisfy the *L. major* FUT1 requirement (Murta et al., 2009).

**FIGURE 10.**
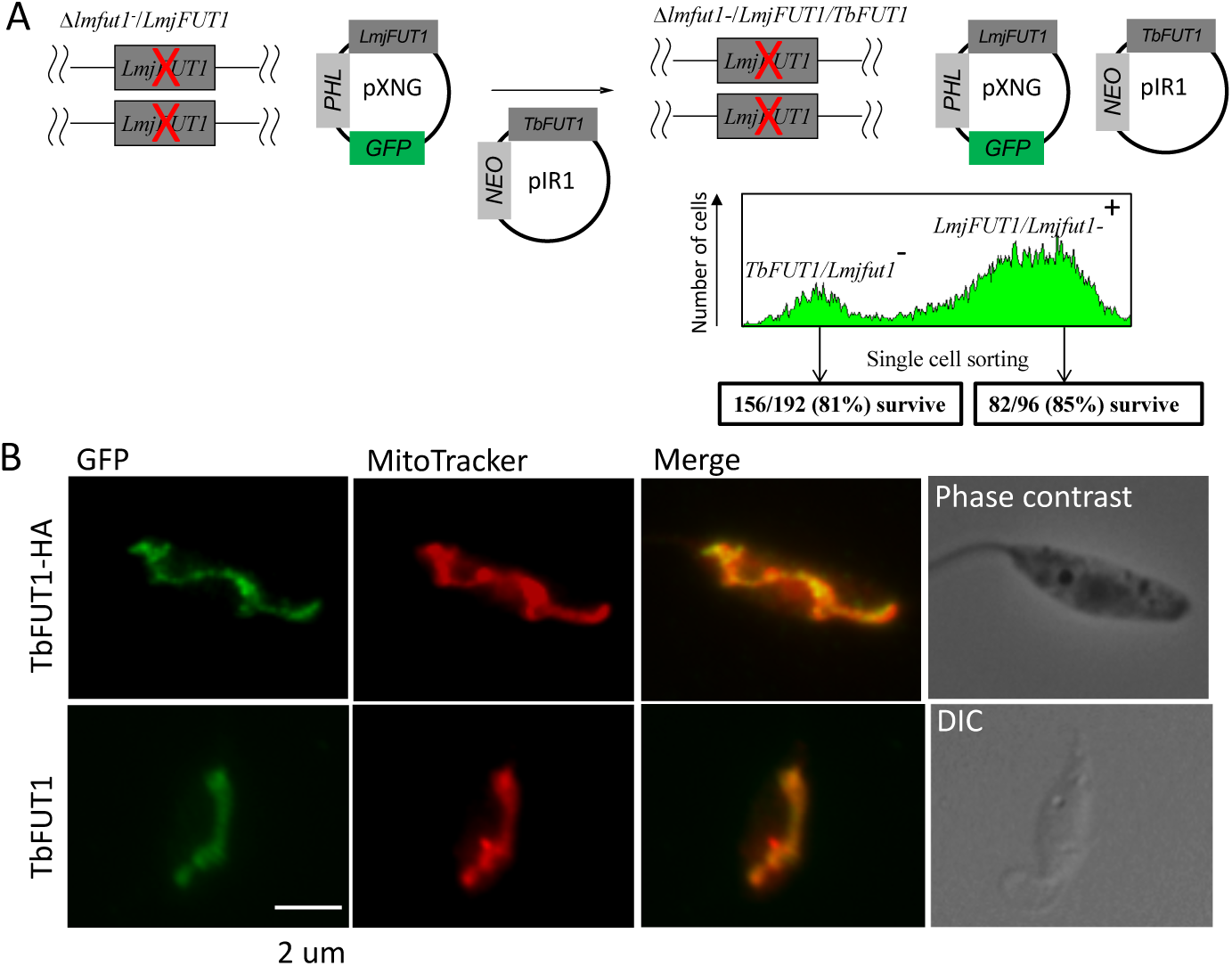
TbFUT1 can functionally and spatially replace LmjFUT1. *A*. Outline of the ‘plasmid shuffling’ procedure to replace *LmjFUT1* with *TbFUT1*. An *L. major* homozygous *fut1*-null mutant expressing *LmjFUT1* (from the GFP-positive, episomal pXNGPHLEO vector) was transfected with pIR1NEO-*TbFUT1*(±HA) expressing either WT or HA-tagged TbFUT1 (GFP-negative). Following transfection and growth in the absence of selective drugs, single cells were sorted for GFP expression or loss. A similar yield of colonies was obtained from both populations. *B.* TbFUT1 HA-tagged (top row) or WT (bottom row) was expressed in an *L. majo*r chromosomal *fut1*-mutant background and analysed by indirect immunofluorescence using anti-HA or anti-TbFUT1 antibodies. Mitotracker red was used as a mitochondrial marker. Scale bar: 2 μm.

Immunofluorescence microscopy of progeny *Lmj fut1*-/pIR1NEO-TbFUT1(±HA) transfected parasites with either mouse monoclonal anti-HA antibodies or affinity-purified rabbit polyclonal anti-TbFUT1 antibodies showed that TbFUT1 was faithfully localized in the *Leishmania* mitochondrion (Fig. 10*B*), The Pearson *r* correlation for co-localization of the anti-HA or anti-TbFUT1 signals with Mitotracker red was > 0.98 in both experiments. In a negative control where anti-TbFUT1 antibody was omitted, the fluorescence signal was lost (data not shown). This result establishes conservation of mitochondrial targeting of TbFUT1 when heterologously expressed in *L. major* promastigotes.

## DISCUSSION

The presence and essentiality of the nucleotide sugar donor GDP-Fuc in BSF and PCF trypanosomes led us to search for putative FUTs in the parasite genome. Only one gene (*TbFUT1*; Tb927.9.3600), belonging to the CAZy GT11 family, was found and phylogenetic analyses revealed that one, or two in the case of *T. cruzi*, orthologues could be found in the genomes of other kinetoplastids. These putative kinetoplastid FUTs form a distinct clade within the GT11 FUT superfamily and are distinct from the GT10 FUTs found in *T. cruzi, L. major* and related parasites, which are absent in *T. brucei*. Consistent with TbFUT1 being the only enzyme likely to utilise the essential metabolite GDP-Fuc, we found that TbFUT1 is also essential to both BSF and PCF parasites. We were able to express TbFUT1 in *E. coli* and the recombinant enzyme was used to demonstrate its activity as a GDP-Fuc : βGal α-1,2 fucosyltransferase with an apparent preference for a Galβ1,3GlcNAcβ1-O-R acceptor motif out of the acceptor substrates investigated.

The highly unusual result was the localization of TbFUT1 to the parasite mitochondrion, using an affinity-purified antibody raised against native TbFUT1 as well as C-terminal epitope tagging. Although in recent years fucosylation has been described in the nucleus and cytosol of protists and plants (Bandini et al., 2016; Rahman et al., 2016; Van Der Wel et al., 2002; Zentella et al., 2017), we are unaware of any other examples of mitochondrial FUTs in any organism. Mitochondrial glycosylation in general is poorly understood. The only other known example of a mitochondrial-localized glycosyltransferase is mitochondrial *O*-GlcNAc transferase (mOGT) (Bond and Hanover, 2015). Mitochondria of rat cardiomyocytes are also positive for OGA and express a UDP-GlcNAc transporter on their outer membrane, indicating all of the molecular components required for this cycling post-translational modification are present in the organelle (Banerjee et al., 2015). Interestingly, disruption of mOGT in HeLa cell mitochondria also leads to mitochondrial disfunction. While there are no (m)OGT orthologues in kinetoplastids, these observations highlight some of the challenges inherent for a mitochondrial-localised glycosyltransferase. Firstly, for TbFUT1 to be active, GDP-Fuc needs to be imported into the mitochondrion, suggesting the presence of an uncharacterized mitochondrial GDP-sugar transporter. Secondly, TbFUT1 appears to be an α-1,2-FUT that decorates glycans terminating in Galβ1,3GlcNAc, suggesting either that additional uncharacterized glycosyltransferases and nucleotide sugar transporters may be present in the parasite mitochondrion, or that the glycoconjugate substrate is assembled in the cytoplasm or secretory pathway and then somehow imported into the mitochondrion to be modified by TbFUT1. Experiments to resolve these options will be undertaken, as will further experiments to try to find the protein, lipid and/or other acceptor substrates of TbFUT1. The latter may then provide clues as to why TbFUT1 is essential for mitochondrial function and parasite growth. Several attempts to identify TbFUT1 substrates have failed so far, including fucose-specific lectin blotting and pull-downs, [^3^H]fucose labelling of parasites transfected with GDP-Fuc salvage pathway enzymes and LC-MS/MS precursor ion and neutral-loss scanning methods. Although the significance is unclear, it is interesting to note that procyclic TbFUT1 has been shown to be under circadian regulation (Rijo-Ferreira et al., 2017).

In conclusion, TbFUT1 is an essential, mitochondrial *α*-1,2-FUT with orthologues throughout the kinetoplastida. As described in (Guo et al. 2021), the activity and mitochondrial localization of the *L. major* homologue are both also required for parasite viability, whereas other putative *L. major* FUTs targeted to the secretory pathway are dispensable. Although no data are available so far on the enzymes from *T. cruzi* or other members of this group, these initial results suggest the intriguing possibility of an essential, conserved mitochondrial fucosylation pathway in kinetoplastids that might be exploitable as a common drug target.

## MATERIAL & METHODS

### Parasite strains

*T. brucei* procyclic form (strain 427, clone 29.13) and bloodstream form (strain 427, variant MITaT 1.2) were used in these experiments. Both strains are stably expressing T7 polymerase and tetracycline repressor protein under G418 (bloodstream) or G418 and hygromycin (procyclic) selection (Wirtz et al., 1999). Procyclic form *T. brucei* was cultured in SDM-79 medium (Brun and Schonenberger, 1979) containing 15% fetal bovine serum and GlutaMAX™ at 28°C. Bloodstream form *T. brucei* was grown in HMI-9t at 37°C, 5% CO_2_. Induction was performed at 0.5 μg/ml tetracycline in both forms.

### BLASTp searches

The *T. brucei, T. cruzi* and *L. major* predicted proteins (from the GeneDB database, http://www.genedb.org) were searched for putative fucosyltransferases using the BLASTp search algorithm (Altschul et al., 1997). The query input sequences are listed in supplemental Table S1. Protein sequence multiple alignments were assembled using Clustal*Ω* (Sievers et al., 2011) and Jalview (Waterhouse et al., 2009).

### Cloning, protein expression and purification of TbFUT1

The open reading frame (ORF) was amplified by PCR from *T. brucei* strain 427 genomic DNA and cloned into the N-terminal GST fusion vector pGEX-6P-1, modified to contain the PreScission Protease site (kind gift of Prof. Daan Van Aalten) using primers P1 and P2 (Table S2). Either the resulting pGEX6P1-GST-PP-TbFUT1 or pGEX6P1-GST-PP were transformed into BL21 (DE3) *E. coli* strain. Recombinant protein expression was induced with 0.1 mM isopropyl-β-D-thiogalactopyranoside (IPTG) and carried out at 16°C for 16 h. Prior to lysis by French press, cells were incubated for 30 min on ice in 50 mM Tris-HCl, 0.15 M NaCl, 1 mM DTT, pH 7.3 (Buffer A) with EDTA-free Complete Protease Inhibitors Tablet (Roche) and 1 mg/ml lysozyme. The soluble fraction, obtained by centrifugation at 17000 x g, 4°C for 30 min, was incubated 2 h at 4°C with Glutathione Sepharose Fast Flow beads that had been pre-equilibrated in Buffer A. The mixture was loaded into a disposable column and the beads washed first with 50 mM Tris-HCl, 0.25 M NaCl, 1 mM DTT, pH 7.3 (Buffer B), then with 50 mM Tris-HCl, 0.15 M NaCl, 0.1 % sodium deoxycholate, 1 mM DTT, pH 7.3 (Buffer C), before being re-equilibrated in Buffer B. GST-TbFUT1 or GST were eluted in 50 mM Tris-HCl, 0.15 M NaCl, 10 mM reduced glutathione pH 8.0 (Buffer E). The yield of recombinant protein was estimated by Bradford assay and the degree of purification by SDS-PAGE analysis. Recombinant protein identification by peptide mass fingerprinting was performed by the Proteomic and Mass Spectrometry facility, School of Life Sciences, University of Dundee.

### Fucosyltransferase activity assays

Aliquots of 2 μg of affinity purified GST-TbFUT1 were incubated with 1 μCi GDP[^3^H]Fuc (American Radiochemicals), 1 mM acceptor in 50 mM Tris-HCl, 25 mM KCl, 5 mM MgCl_2_, 5 mM MnCl_2_, pH 7.2 for 2 h at 37°C. The acceptors tested (Table 1) were purchased from Sigma, Dextra Laboratories or Toronto Research Chemicals. To study the dependency on divalent cations, MgCl_2_ and MnCl_2_ were removed from the buffer and a formulation with 10 mM EDTA was also tested. Reactions were stopped by cooling on ice then desalted on mixed bed as follow: 100 μl each Chelex100 (Na^+^) over Dowex AG50 (H^+^) over Dowex AG3 (OH^-^) over QAE-Sepharose A25 (OH^-^). The flow through and the elutions (4 x 400 μl of water) were combined. About 5% of the desalted reactions were counted at a LS 6500 scintillation counter (Beckmann). The remaining material was lyophilized for further analyses.

### HPTLC analysis

Reaction products and standards were dissolved in 20% 1-propanol and separated on a 10 cm HPTLC Si-60 plates (Merck) using 1-propanol:acetone:water 9:6:4 (v:v:v) as mobile phase. Non-radiolabelled sugars were visualized by orcinol/ H_2_SO_4_ staining. In the case of radiolabelled products, the HPTLC plates were sprayed with En^3^hance^®^ (PerkinElmer) and visualized by fluorography.

### Large scale TbFUT1 assay and product purification

Acceptor (5 mM Lacto-N-biose-β-O-methyl) and donor (2.5 mM GDP-Fuc) were incubated with 8 μg affinity purified GST-TbFUT1 in 20 mM Tris-HCl, 25 mM KCl, pH 7.2 at 37°C for 24 h. The reaction products were desalted on a mixed-bed column and lyophilized (see above). The trisaccharide product was then isolated by normal phase liquid chromatography. The reactions were dissolved in 95% acetonitrile (ACN) and loaded on a Supelco SUPELCOSYL™ LC NH_2_ HPLC column (7.5 cm x 3 mm, 3μm) using an Agilent 1120 Compact LC system. The purification was performed at 40°C with a 0.3 ml/min flow rate. The column was equilibrated in 100% Buffer A (95 % ACN, 5% H_2_O) for 5 min, before applying a first gradient from 0-35 % Buffer B (95% H_2_O, 5 % ACN, 15 mM ammonium acetate) over 25 min followed by a second gradient from 35-80 % Buffer B over 20 min. The column was then re-equilibrated in 100% Buffer A for 10 min. About 1.5 % of each fraction was potted onto a silica HPTLC plate and stained with orcinol/ H_2_SO_4_ staining to identify the sugar-containing fractions. An aliquot from each orcinol-positive fraction was then analysed by HPTLC and the ones containing the putative trisaccharide product were pooled and freeze-dried.

### Permethylation, ESI-MS analysis and GC-MC methylation linkage analysis

Purified TbFUT1 reaction product was dried and permethylated by the sodium hydroxide method as described in Ferguson, 1992. Aliquots were used for ESI-MS and the remainder was subjected to acid hydrolysis followed by NaB^2^H_4_ reduction and acetylation (Ferguson, 1992). The resulting PMAAs were analysed using an HP6890 GC System equipped with an HP-5 column linked to a 5975C mass spectrometer (Agilent). For ESI-MS and ESI-MS/MS, the permethylated trisaccharide was dried and re-dissolved in 80% ACN containing 0.5 mM sodium acetate and loaded into gold-plated nanotips (Waters) for direct infusion. The Q-Star XL mass spectrometer equipped with Analyst software (Applied Biosystems) was operated in positive ion mode using an ion spray voltage of 900 V and a collision voltage of 50-60 V in MS/MS mode.

### NMR

The purified TbFUT1 reaction product was exchanged in ^2^H_2_O by freeze-drying and analysed by one-dimensional ^1^H-NMR and two-dimensional ^1^H-ROESY (Rotating frame Overhouser Effect SpectroscopY). All spectra were acquired on a Bruker Avance spectrometer operating at 500 MHz with a probe temperature of 293°K.

### Generation of *TbFUT1* conditional null mutants

About 500 bp of the 5’ and 3’ untranslated regions (UTRs) immediately flanking the *TbFUT1* ORF were amplified from *T. brucei* 427 genomic DNA (gDNA) using primers P3/P5 and P4/P6 (Table S2) and linked together by PCR and the final product was ligated into pGEM-5Zf(+). Antibiotic resistance cassettes were cloned into the *Hind*III/*BamH*I restriction sites between the two UTRs to generate constructs either containing puromycin acetyltransferase (*PAC*) or blastacidin S deamidase (*BSD*). To allow cloning of the antibiotic resistance cassettes using *Hind*III/*Bam*HI, a *BamH*I site present in the 5’-UTR was first mutated to gg*c*tcc using the QuickChange Site-Directed Mutagenesis Kit (Stratagene) and primers P7/P8. In addition, a hygromycin (HYG) based *TbFUT1* gene replacement cassette was generated with longer UTRs (1.25 kb) amplified using primers P9/P10 (5’ UTRs) and P11/P12 (3’UTRs). To avoid the endogenous 5’ UTR site, first the 3’UTR was introduced via *BamH*I/*Sac*I then the 5’UTR via *Spe*I/*Hind*III. The tetracycline-inducible ectopic copy construct was generated by amplifying the *TbFUT1* ORF from *T. brucei* gDNA (primers P13/P14) and cloning the resulting PCR product into pLEW100. Additionally, pLEW100XM was generated which allowed universal tagging of a protein of interest with a C-terminal MYC_3_ tag (see details below). Linearized DNA was used to transform the parasites as previously described (Burkard et al., 2007; Güther et al., 2006; Wirtz et al., 1999). The genotype of the transformed parasites was verified by Southern blot as detailed below.

### Southern blotting

Genomic DNA was prepared from 5×10^7^ or 1×10^8^ cells using DNAzol (Helena Biosciences) and digested with 0.1 mg/ml RNAse I. The gDNA (5 μg/lane) was digested with the appropriate restriction endonucleases and probed with the ORFs of *TbFUT1*, puromycin acetyltransferase gene (*PAC*), hygromycin B phosphotranferase (*HYG*) or blastacidin S deamidase gene (*BSD*) generated using the PCR DIG Probe Synthesis Kit (Roche). The blot was developed according to the manufacturer’s instructions.

### Northern blotting

Total RNA was prepared from 5×10^6^-1×10^7^ cells using the RNeasy MIDI Kit (Qiagen) according to manufacturer’s instructions. The RNA was separated on a 2% agarose-formaldehyde gel, blotted and detected using the Northern Starter Kit (Roche). Probes were designed based on the DIG RNA Labelling Kit T7 (Roche) and *TbFUT1* and alpha-tubulin (Tb427.01.2340) templates were amplified from *T. brucei* bloodstream form gDNA using primers P17/P18 and P19/P20 (Table S2), respectively. Total RNA and DIG labelled probes were quality checked by capillary electrophoresis on an Agilent BioAnalyzer 2100.

### Generation and expression of epitope-tagged TbFUT1 constructs

*TbFUT1* was introduced in two different sites of pLEW100HXM to yield HA_3_-TbFUT1 and HA_3_-TbFUT1-MYC_3_. To modify the pLEW100 vector, we inserted a 99 bp adapter oligo encoding for the following sequence *Hind*III/*Nde*I/*Asc*I/*BbvC*I/*Xba*I/*BstB*I/*Xho*I/<u>HA</u>**/***Pac*I/*BamH*I, where <u>HA</u> stands for a single HA tag (O1 and O2 in Table S2). The HA tag was then replaced with a MYC_3_ tag amplified from a pMOTag43M plasmid, creating the following multiple cloning site: *Hind*III/*Nde*I/*Asc*I/*BbvC*I/*Xba*I/*BstB*I/*Xho*I/<u>MYC_3_</u>**/***Pac*I/*BamH*I. This vector was given the name pLEW100XM. *TbFUT1* was amplified from a plasmid source using primers P15/P16 and inserted via *Hind*III/*Xho*I. pLEW100XM was then used to introduce an HA_3_ adapter which would allow N-terminal tagging of a protein of choice. The adapter consisted of the O3/O4 primer pair (Table S2). The primer pair was fused and inserted into *Hind*III/*Asc*I restriction sites of pLEW100XM creating the following multiple cloning site: *Hind*III/**HA_3_**/*Asc*I/*BbvC*I/*Xba*I/*BstB*I/*Xho*I/**MYC_3_/***Pac*I/*BamH*I. This plasmid was termed pLEW100HXM. TbFUT1 was amplified using primers P21/P22 for HA_3_-TbFUT1 and P23/P16 for HA_3_-TbFUT1-MYC_3_ (Table S2). The two plasmids were purified and electroporated into BSF cells as described above. The generation of the TbFUT1-MYC_3_ cell line is described above as it was used as parental strain for the BSF *TbFUT1* cKO cell line.

### Preparation of anti-FUT1 antibody

The *Nde*I site within the *TbFUT1* sequence was eliminated using the QuickChange Lightning Site-Directed Mutagenesis Kit (Agilent) and primers P24/P25. The resulting mutagenized ORF was then amplified with primers P26/P27 and cloned into pET15b resulting in an N-terminally truncated construct encoding Δ_32_TbFUT1 fused to an N-terminal hexahistidine tag (HIS_6_) with PreScission plus (PP) protease cleavage site. *E. coli* BL21 gold (DE3) cells were allowed to express His_6_-PP-TbFUT1 overnight at 25°C in auto-inducing media (5052-NPS-MgSO_4_). Cells were harvested by centrifugation and incubated for on ice in 50 mM Tris-HCl pH 7.3, 0.15 M NaCl, 1 mM DTT (Buffer A) with EDTA-free Complete Protease Inhibitors Tablet (Roche), before being disrupted in a French Press. Since all of the protein was insoluble within inclusion bodies a refolding procedure was applied. The cell lysate was spun down for 20 min at 20,000x g and the pellet was washed in ice cold PBS with EDTA-free Complete Protease Inhibitors Tablet (Roche). Cell debris layering the inclusion body pellet was scratched off and the pellet washed again. This process was repeated 3 times. The inclusion body pellet was washed with PBS + 0.5% TX-100 and sonicated on ice. After digestion of residual DNA and RNA, the pellet was washed with PBS + 0.1% Tween-20, sonicated, and washed in TBS. The washed inclusion bodies were resuspended in 8 M UREA/ TBS and incubated overnight at 37°C, 200 rpm. Insoluble material was spun down and the supernatant slowly diluted down under agitation to 2 M UREA with TBS at room temperature. After centrifugation the supernatant concentrated to 2 mg/mL in a 30 kDa cut-off VivaSpin concentrator. Re-solubilised protein (2 mg) was sent to DC Biosciences ^LTD^ for production of polyclonal rabbit antiserum.

The IgG fraction from the hyperimmunized rabbit serum was purified on 2x 1 mL HiTrap™ Protein G HP columns (GE Healthcare). And then affinity purified on GST-TbFUT1 fusion protein coupled to a HiTrap NHS-Activated HP column (GE Healthcare). Finally, the IgG fraction was adsorbed against PCF *TbFUT1* cKO cell lysates prepared on day 9 day without tetracycline, a time point at which they are virtually free of TbFUT1. The cell lysate was prepared by sonication of 2×10^8^ cells on ice in 100 mM sodium phosphate buffer pH 7 including EDTA-free Complete Protease Inhibitors (Roche) and 0.01% TX-100. After addition of purified TbFUT1-specific IgGs to the cell lysate, the solution was incubated rotating overnight at 4°C, concentrated in a VivaSpin 30 kD cut-off filter and stored in 50% glycerol at −20°C. Final estimated rabbit anti-TbFUT1 IgG concentration was 50 µg/mL.

### Immunofluorescence microscopy

Late log phase *T. brucei* bloodstream form or procyclic cells were fixed in 4% PFA/PBS in solution at a concentration of 5×10^6^ cells/mL. When using Mitotracker^TM^ Red CMX Ros cell cultures were spiked with a 25 nM concentration over 20 minutes, before harvesting. Fixed parasites were permeabilized in 0.5% TX-100 in PBS for 20 min at room temperature, before blocking in 5% fish skin gelatine (FSG) containing 0.1% TX-100 and 10% normal goat serum. Alternatively, procyclic cells were permeabilised in 0.1% TX-100 in PBS for 10 min and blocked in 5% FSG containing 0.05% TX-100 and 10% normal goat serum. Primary antibodies incubations were performed in 1% FSG, 0.1% TX-100 in PBS. The following primary antibodies were used in 1:1000 dilutions: rabbit anti-TbFUT1 IgG (see above), rabbit anti-GRASP IgG (gift from Prof. Graham Warren, University College London), rabbit anti-ATPase IgG (gift from Prof. David Horn, University of Dundee), mouse anti-HA IgM (Sigma), rabbit anti-HA IgG (Invitrogen), mouse anti-MYC IgM (Millipore). Anti-FAZ was used at 1:2 and anti-PFR at 1:10 (both antibodies were kind gifts of Prof. Keith Gull, University of Oxford). After washing, the coverslips were incubated with a 1:500 dilution of respective secondary Alexa Fluor^®^ goat IgG, like anti-mouse 488 and 594, as well as anti-rabbit 488 and 594 (Life Technologies). Coverslips were mounted on glass slides with Prolong® Gold DAPI antifade reagent (Invitrogen). Microscopy was performed on a DeltaVision Spectris microscope (GE Healthcare) and images were processed using Softworx. For Figure S10, microscopy was performed on a Zeiss LSM 700 confocal microscope. All images were further processed using Fiji (Schindelin et al., 2012).

### *Leishmania major* complementation by ‘plasmid shuffling’

For the plasmid swap experiments described in Fig. 10, the *TbFUT1* ORF was amplified by PCR and inserted into the expression vector pIR1NEO yielding pIR1NEO-TbFUT1 (B7148). *L. major* parasites were grown as promastigotes in M199 medium, transfected and plated on selective media for clonal lines as described previously (Guo et al., 2017). Plasmid ‘shuffling’ to swap an episomal *LmjFUT1*-expressing construct with the ones expressing-TbFUT1(± HA tag) was performed as described in the text and Fig. 10A, following procedures described in (Guo et al., 2017).

## ACKNOWLEDGEMENTS

The authors would like to thank Gina MacKay and Art Crossman (University of Dundee) for performing the NMR experiment and helping with the data analysis. We would also like to thank Alan R. Prescott (Division of Cell Signalling and Immunology, University of Dundee) for his generous help with the confocal microscopy and Martin Kierans for preparing the samples for scanning electron microscopy. The authors are also grateful to Keith Gull (University of Oxford), Graham Warren (University College London), Daan van Aalten and David Horn (University of Dundee) for providing reagents. This work was supported by a Wellcome Trust Investigator Award (101842) to MAJF, University of Dundee/BBSRC PhD studentship to GB, NIH Grant R01-AI31078 to SMB and postdoctoral fellowship to HG.

## AUTHOR CONTRIBUTIONS

G.B., S.D., S. M. B. and M.A.J.F. designed research; G.B., S.D., H. G. and M.L.S.G. performed research; G.B., S.D., A. M., M.L.S.G., S. M.B. and M.A.J.F. analysed data; G.B., S.D., S.M.B. and M.A.J.F. wrote the paper.

## Supplementary Information for

### Supplementary Methods

#### Recovery of radiolabelled products

The reaction products from the TbFUT1 activity assays using Lac, LNB and LacNAc as acceptors were separated on a 20 cm HPTLC Si-60 plate. The radiolabelled products were localized on the TLC plate using an AR-2000 (BioScan) plate reader, the corresponding areas were wetted with mobile phase and the silica scraped and transferred to microcentrifuge tubes. The radiolabelled materials were extracted from the solid phase by incubating with mobile phase and a 5% aliquot counted at the scintillation counter to determine the amount of recovered material.

#### Acid hydrolysis

Aliquots corresponding to 15% of the extracted radiolabelled reaction products were dried in a Speedvac concentrator, re-dissolved in 4 M TFA and incubated at 100°C for 4 h (1). After cooling, the samples were dried on a Speedvac concentrator, washed in water and re-dissolved in 20% 1-propanol to be analysed by HPTLC as described in the main text.

#### *Xanthomonas manihotis* α1,2-fucosidase reaction

Aliquots corresponding to 15% of the extracted radiolabelled reaction products were treated or mock treated with 10 U *X. manihotis* α-1,2-fucosidase (New England Biolabs) at 37°C for 16 h. The reactions were stopped by heating at 100°C for 10 min and desalted on mixed bed columns as described for the activity assay. After freeze-drying, samples were washed with water, re-dissolved in 20% 1-propanol and analysed by HPTLC.

#### Scanning electron microscopy

Samples were prepared and analysed as described in (2). Briefly, PCF *TbFUT1* cKO cells were grown in presence or absence of tetracycline and fixed on days 6 and 12 in SDM-79 with 2.5% gluteraldehyde and the samples were further prepared by the Dundee Imaging Facility, School of Life Sciences. Cells were collected on 1 μM Shandon Nuclepore membrane filters, rinsed twice in 0.1 M PIPES buffer pH 7.2 and treated with 0.2% osmium tetroxide overnight. Samples were then washed in water and in a gradient of ethanol solutions to dehydrate them. After critical point drying on a BALTec critical point dryer D30, the samples were mounted on aluminium stubs that were then coated with 40 nM gold/palladium. The stubs were examined in a Philips XL30 environmental scanning electron microscope operating at an accelerating voltage of 15 kV.

#### Cell volume determination

The average cell volume was obtained by analysing PCF *TbFUT1* cKO and *TbGMD* cKO on a CASY^®^ Cell Counter + Analyser system. Procyclic form cultures grown in the presence of absence of tetracycline were diluted 1:10 in 1xPBS before being diluted in the CASY-ton solution for measurements.

#### Sedimentation assay

The assay was adapted from (3). Briefly, about 5×10^6^ *T. brucei* PCF *TbFUT1* cKO or *TbGMD* cKO cells were resuspended in 1 ml of SDM-79 medium and transferred into a 2 ml polystyrene cuvette (Sarstedt). The cuvettes were incubated at 28°C (CPS-controller, Shimadzu) in a UV-1601 Spectrophotometer (Shimadzu) and the optical density at 600 nm measured every 30 min. SDM-79 medium was used as a blank.

---

**Table S1.** Summary of BLASTp searches for putative fucosyltransferases in the three kinetoplastids genomes.

| CAZy family | Linkage | Query input | BLASTp results |  |  |
| --- | --- | --- | --- | --- | --- |
|  |  |  | *T. brucei* | *T. cruzi* | *L. major* |
| GT10 | α1,3/1,4 | CAC38049.1 (*A. thaliana*),<br>AAA92977.1 (*HsFUT4*),<br>AAH36037.1 (*HsFUT11*) | | TcCLB.509233.40,<br>TcCLB.509437.80 | LmjF.2.0330 |
| GT11 | α1,2(α1,3) | AAC24453.1 (*HsFUT2*),<br>NP_001316806.1 (*HsFUT1*) | Tb927.9.3600 | TcCLB.506893.90,<br>TcCLB.508501.240 | LmjF.01.0100 |
| GT23 | α1,6 | BAA19764.1 (*HsFUT8*) | | | |
| GT37 | α1,2 | AAD41092.1 (*A. thaliana*) | | | |
| GT56 | α1,4 | AAK62972.1 (*E.coli*) | | | |
| GT65 | O-Fuc | AAL09576.1 (*HsPOFUT1*) |  |  |  |
| GT68 | O-Fuc | AAK77300.1 (*D. melanogaster*) |  |  |  |
| GT74 | α1,2/GalT | Q54QG0.1(*D. discoideum*) | | | |

**Table S2.**
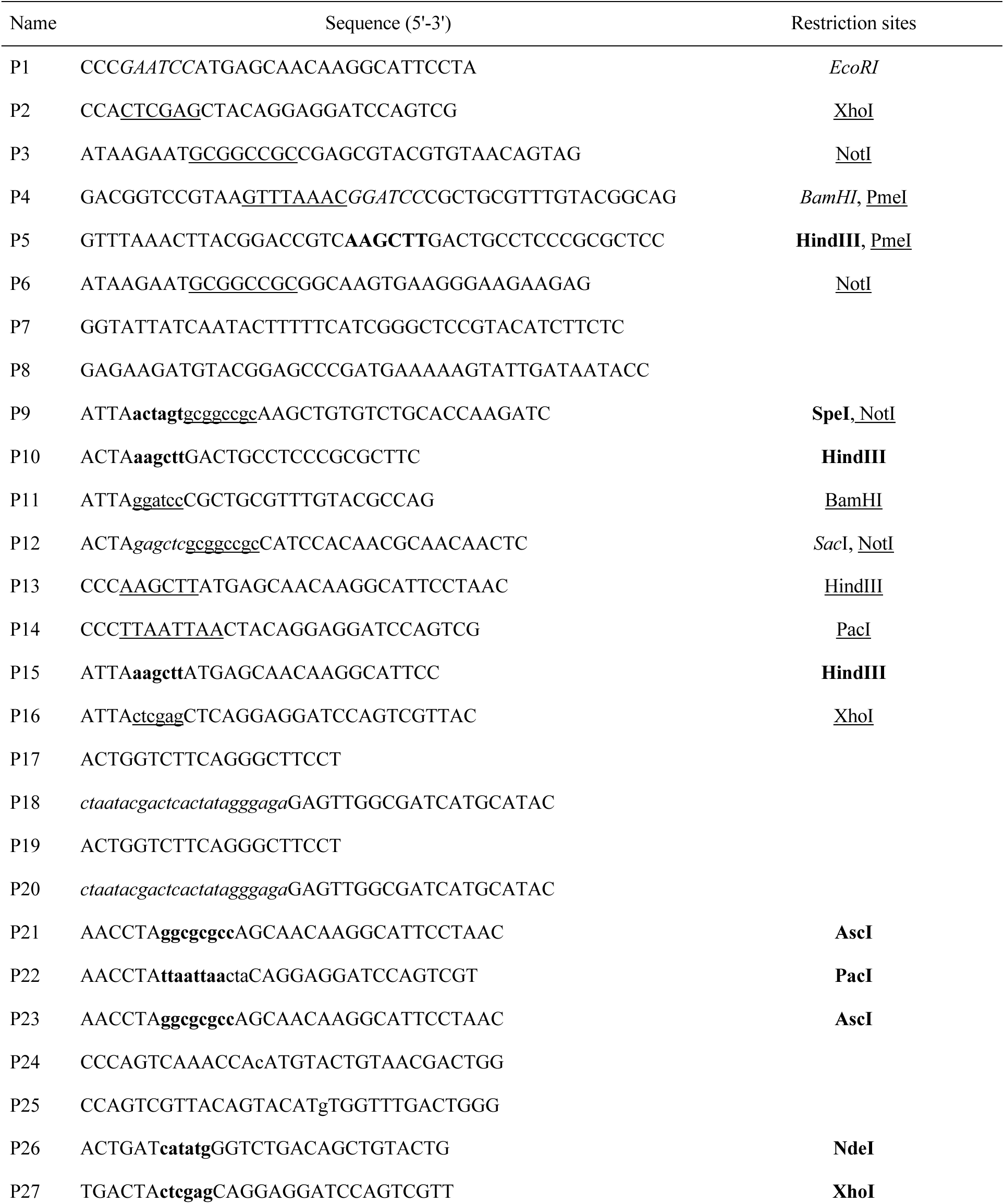
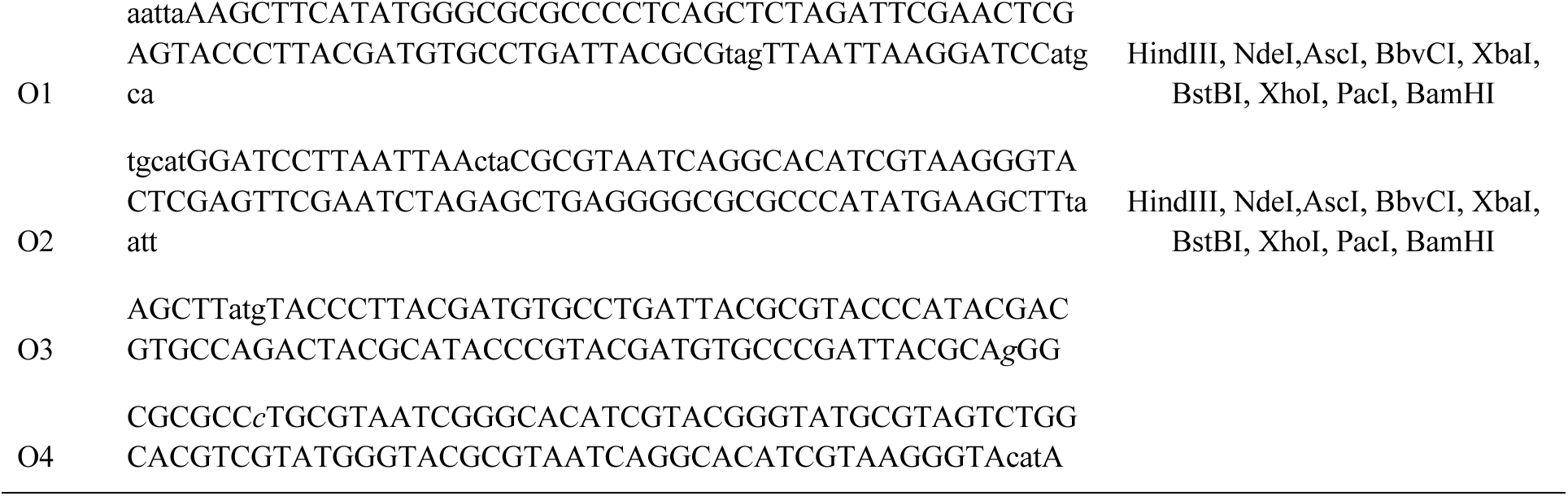
Primers and oligos used in this study

## Supporting Figures S1-S11

**FIGURE S1.**
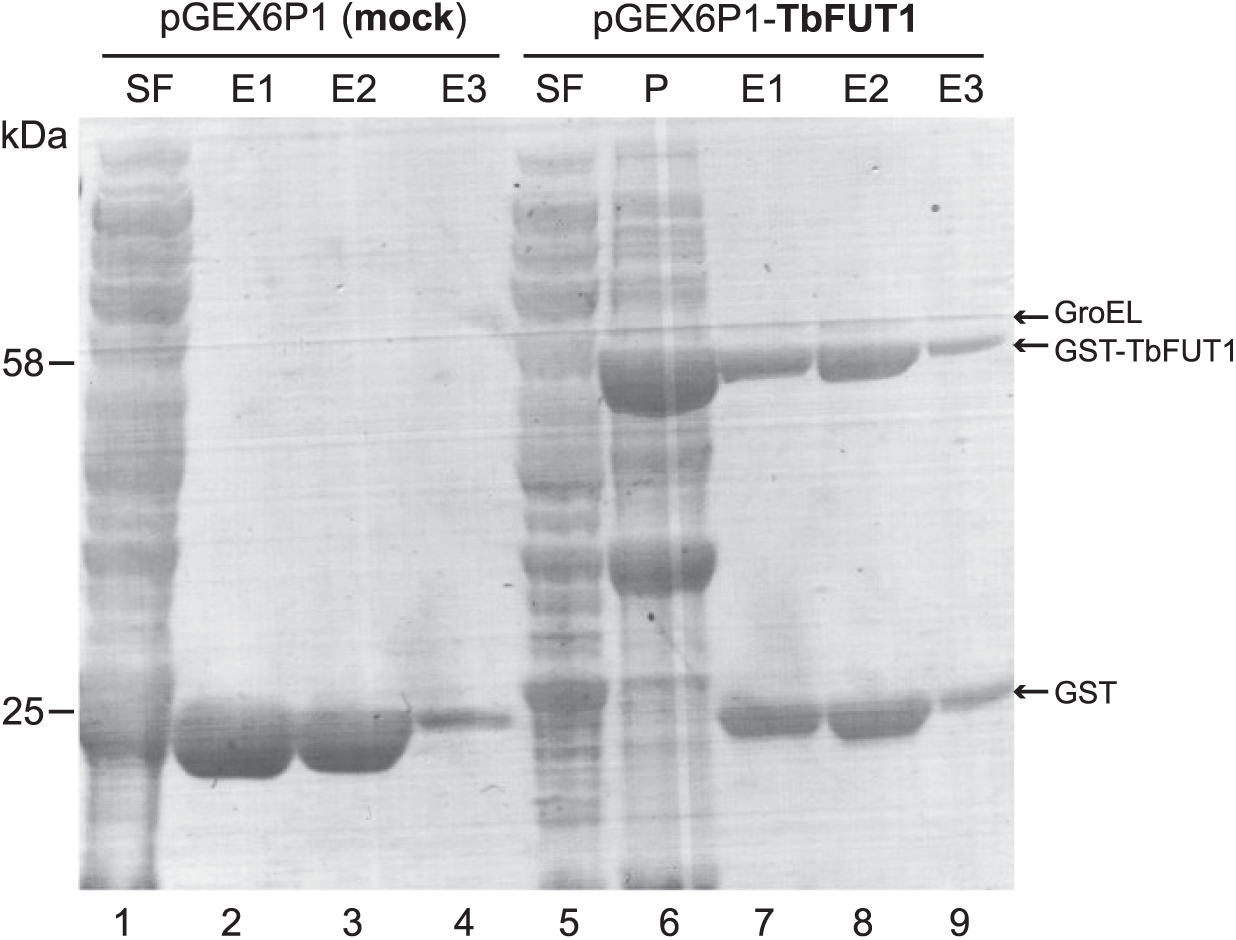
Purification of recombinant GST-TbFUT1. The material encoded by the empty vector (lanes *1-4*) and the GST-TbFUT1 (lanes *5-9*) were expressed and purified following the same protocol (see experimental procedures). Aliquots from the lysis and affinity purification steps were run on a SDS-PAGE and stained with Coomassie. *SF*: soluble fraction. *P*: pellet (insoluble fraction). *E1-E3*: elutions.

**FIGURE S2.**
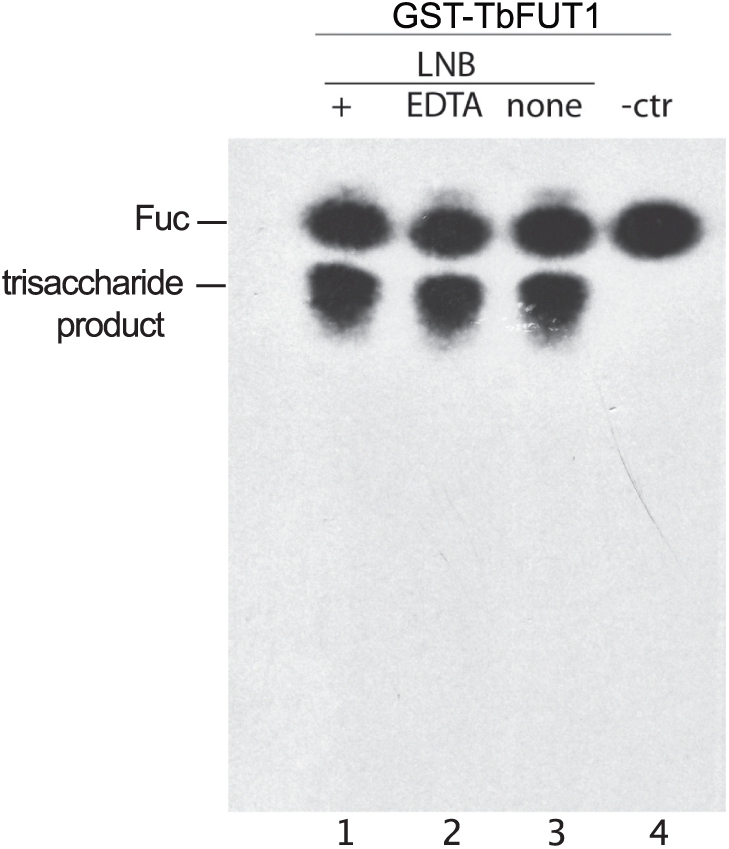
TbFUT1 activity is independent of divalent metal cations. We performed activity assays using lacto-*N*-biose (LNB) as acceptor (see experimental procedures) either using the complete reaction buffer (*+,* lanes *1* and *4*), removing MgCl_2_ and MnCl_2_ from it (*none*, lane 3, 50 mM TrisHCl, 25 mM KCl pH 7.2) or adding EDTA to it (*EDTA*, lane *2*, 50mM TrisHCl, 25 mM KCl, 5 mM EDTA pH 7.2). A negative control missing the LNB acceptor (*- ctr*, lane *4*) was also performed.

**FIGURE S3.**
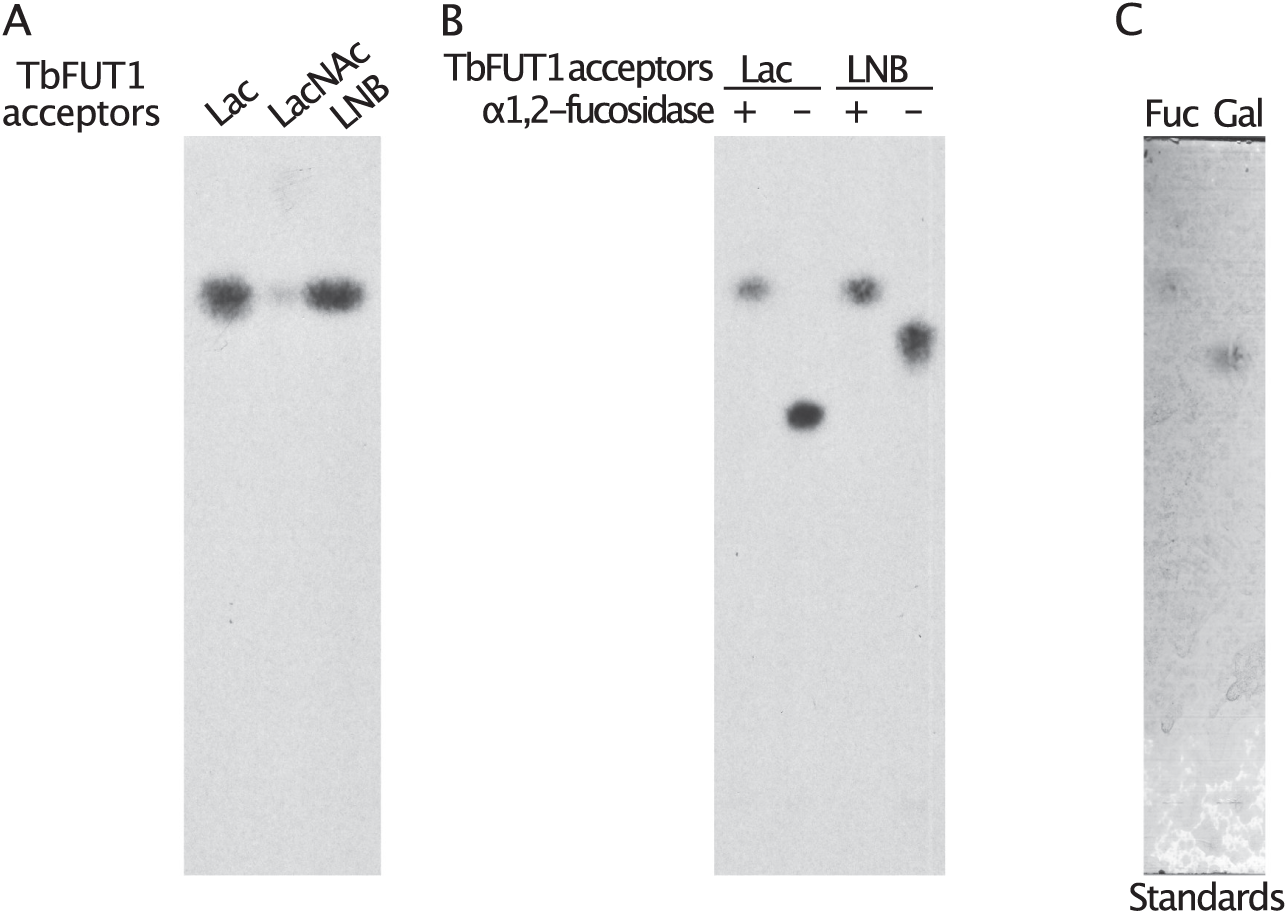
Preliminary characterization of TbFUT1 reaction products. *A*. Labelled products were recovered from the activity assays and treated with 4 M TFA (acid hydrolysis). *B*. Tritiated products, recovered from the TbFUT1 activity assays, were treated (lanes *1* and *3*) with 10 U of *X. manihotis* α-1,2-fucosidase or mock treated (lanes *2* and *4*) by incubating with only the fucosidase buffer. Acid hydrolysis reactions, fucosidase treatments and monosaccharide standards were run on a TLC plate in solvent A. The radiolabelled products were visualized by fluorography and the standards were visualized with orcinol/sulphuric acid (*C*).

**FIGURE S4.**
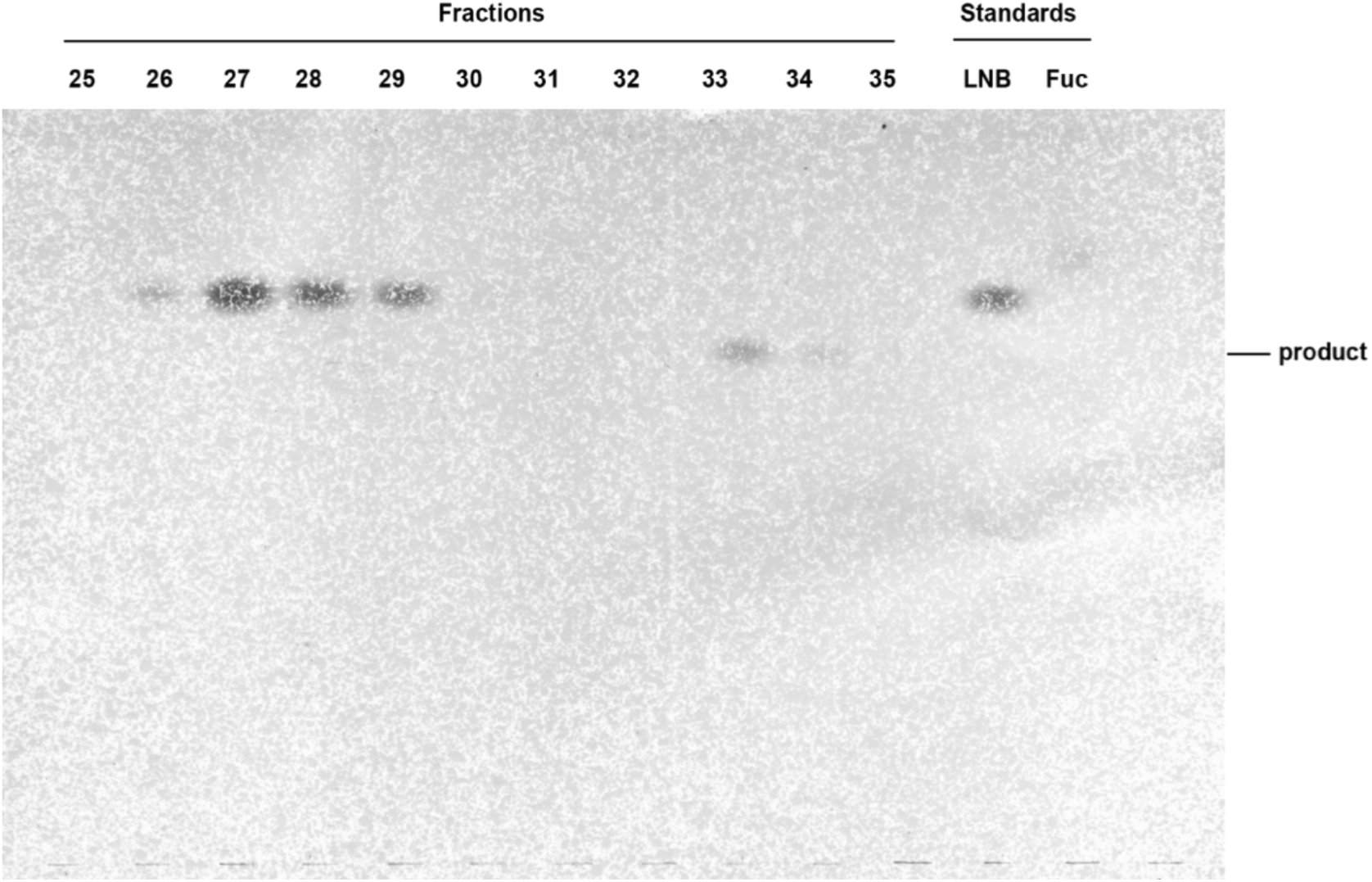
Purification of the TbFUT1 reaction product by normal phase HPLC. An aliquot (2%) from each sugar-containing fraction from the normal phase purification was run on a TLC plate in solvent A and developed with orcinol/sulphuric acid. Standards for the acceptor (LNB) and for Fuc were also analyzed. Fractions containing the reaction product (33 and 34) were pooled for analysis.

**FIGURE S5.**
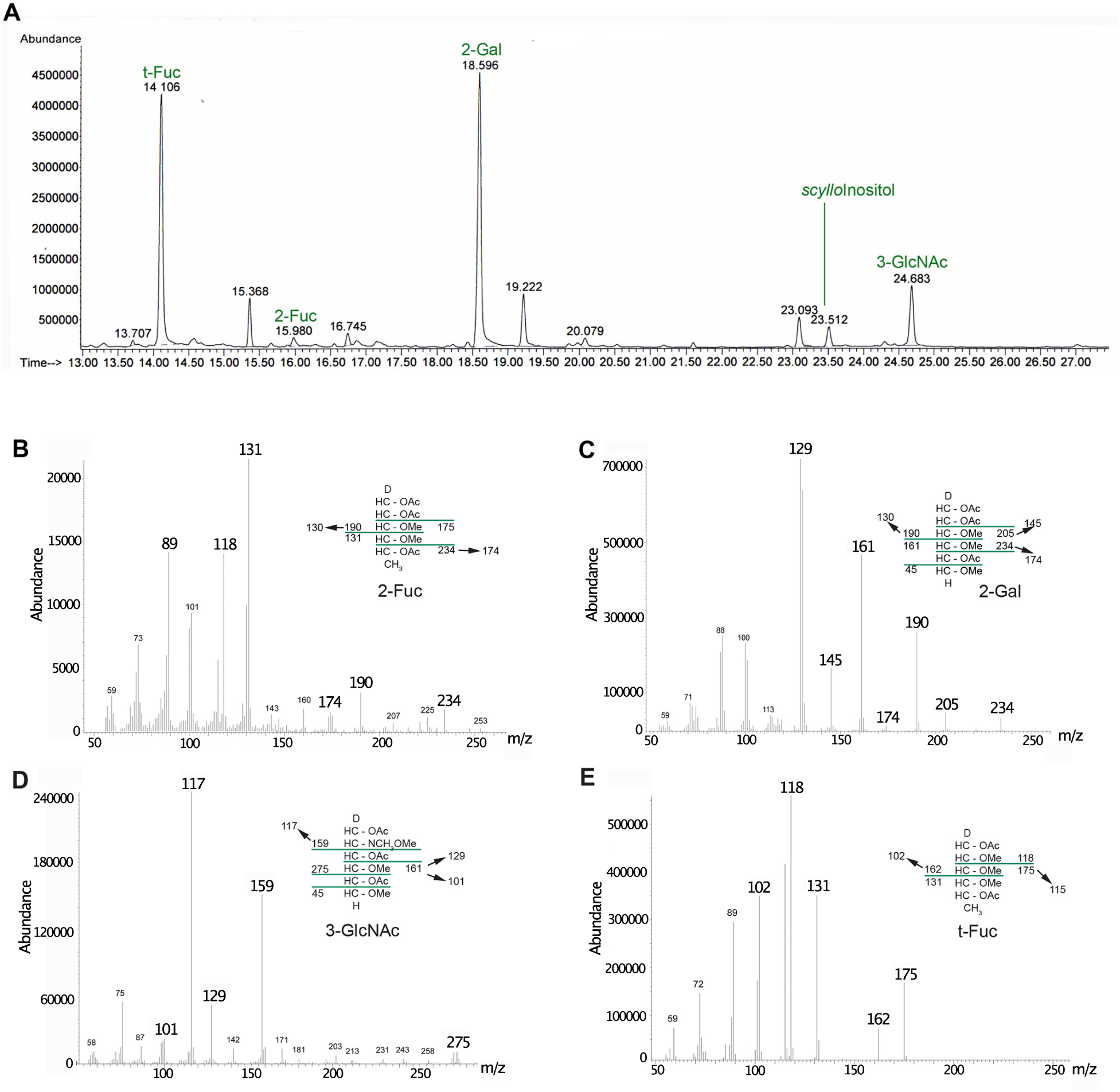
GC-MS methylation linkage analysis of the purified reaction product. *A*. Chromatogram of the PMAAs obtained from the reaction product. The peaks are annotated to reflect their origin in the oligosaccharide: t-Fuc= non-reducing terminal fucose, 2-Gal= 2-*O*-substituted galactose, 3-GlcNAc= 3-*O*-substituted *N*-acetylglucosamine, 2-Fuc=2-*O*-substituted fucose. *B-E*. Electron-impact MS spectra for the peaks at 15.9, 18.6, 24.7 and 14.1 min, respectively. The insets in each panel show the fragmentation patterns.

**FIGURE S6.**
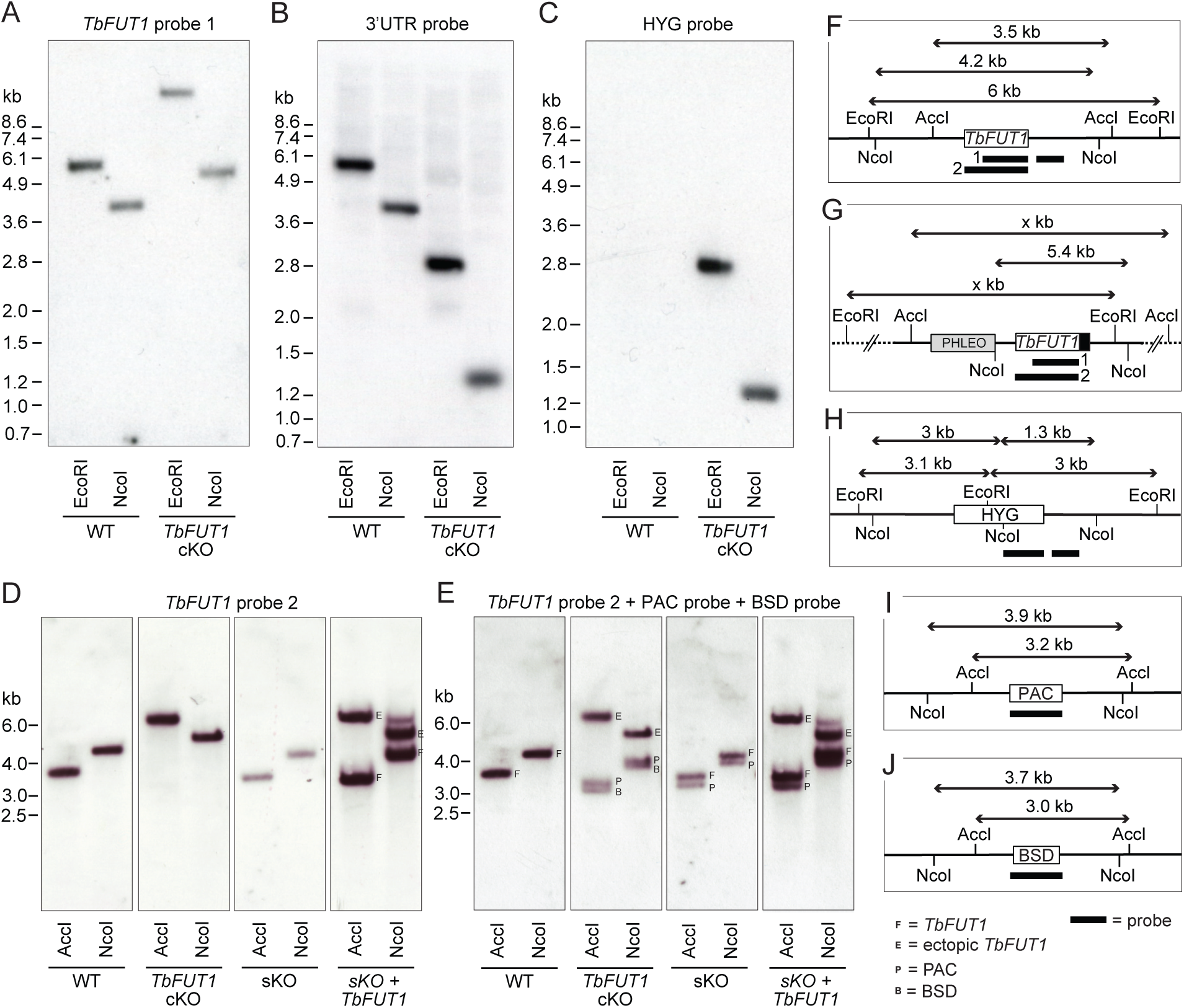
Southern blots of *TbFUT1* conditional null mutants. Bloodstream form gDNA of wildtype (WT) and *TbFUT1* cKO mutant was digested with EcoRI and NcoI, respectively, and three separate blots probed with a DIG-labelled DNA fragment of *TbFUT1* (*A*), 3’UTR (*B*) and HYG (*C*). Similarly, procyclic form WT, single KO (sKO), sKO plus ectopic *TbFUT1 (*sKO *+ TbFUT1)* and *TbFUT1* cKO gDNA was digested with AccI and NcoI, respectively, and one blot consecutively probed with a DIG-labelled DNA fragment of *TbFUT1* (*D*) or PAC and BSD (*E*). A graphic key illustrates the resulting band patterns and fragment sizes expected by probing against the native *TbFUT1* (*F*), the ectopic *TbFUT1* with or without MYC_3_-tag (black box) (*G*) and the three resistance genes HYG (*H*), PAC (*I*) and BSD (*J*).

**FIGURE S7.**
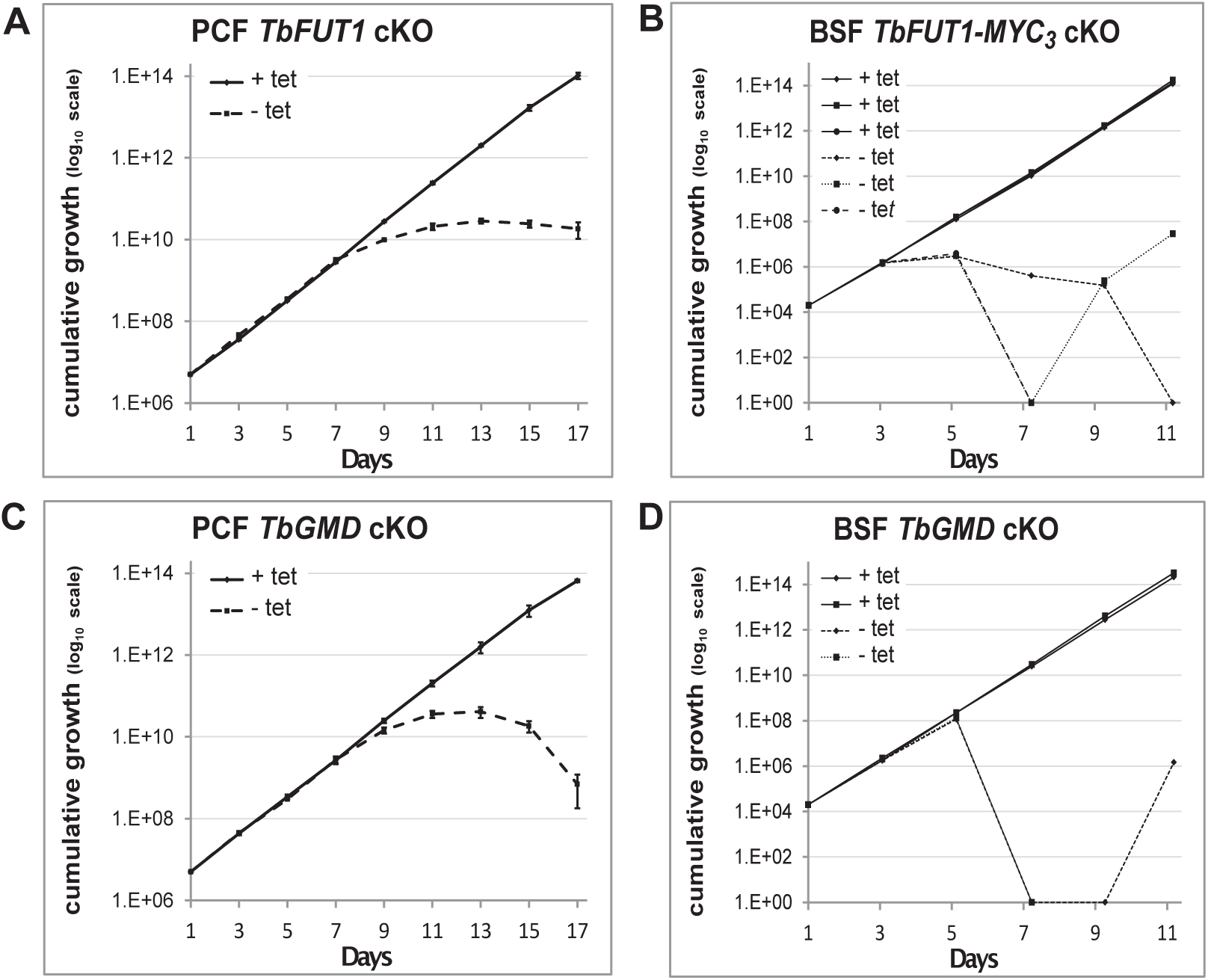
Comparison of *TbFUT1* and *TbGMD* conditional null mutant growth. *TbFUT1* and *Tb GMD* cKO were grown in parallel under permissive (*solid*) or non-permissive (*dotted*) conditions. Cells were split and counted every 2 days in biological duplicates (procyclic form) or triplicates (bloodstream form) using a haemocytometer. Growth is depicted in cumulative curves over several passages of: *A*. Procyclic form *TbFUT1* conditional knockout. *B*. Three different clones of bloodstream form BSF *Tb*FUT1 cKO cells expressing the MYC_3_-tagged ectopic copy. *C*. Procyclic form *TbGMD* cKO cells. *D.* Two different clones of bloodstream form *TbGMD c*KO cells.

**FIGURE S8.**
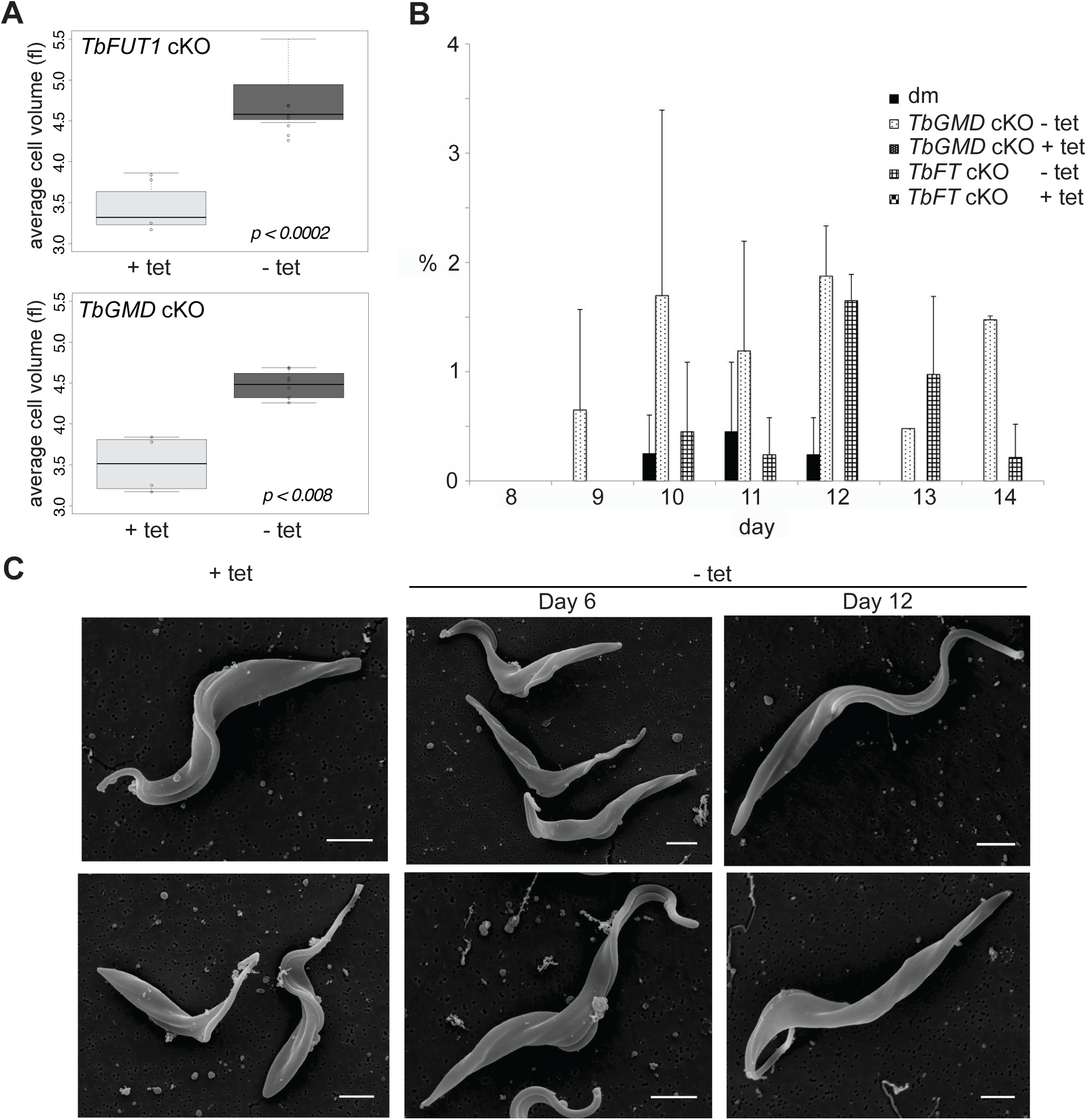
*TbFUT1* and *TbGMD* cKO have increased cell size. *A*. The average cell volume was measured using a Cazy cell counter (day 10 to 13) for both cell lines grown in permissive (+ Tet) and non-permissive (- Tet) conditions. *B*. Number of cells with detached flagella in *TbFUT1* and *TbGMD cKO* cells grown in permissive and non-permissive conditions compared to wild type (dm). The average of two biological replicates is shown. In each replicate 200 cells /cell line were counted using a haemocytometer at an inverted light microscope. *C*. Scanning electron microscopy (SEM) images of PCF *TbFUT1* cKO grown in the presence or absence of tetracycline for 6 and 12 days. No flagellar detachment was observed, but cells grown in non-permissive conditions appear longer. Scale bars: 2 μm.

**FIGURE S9.**
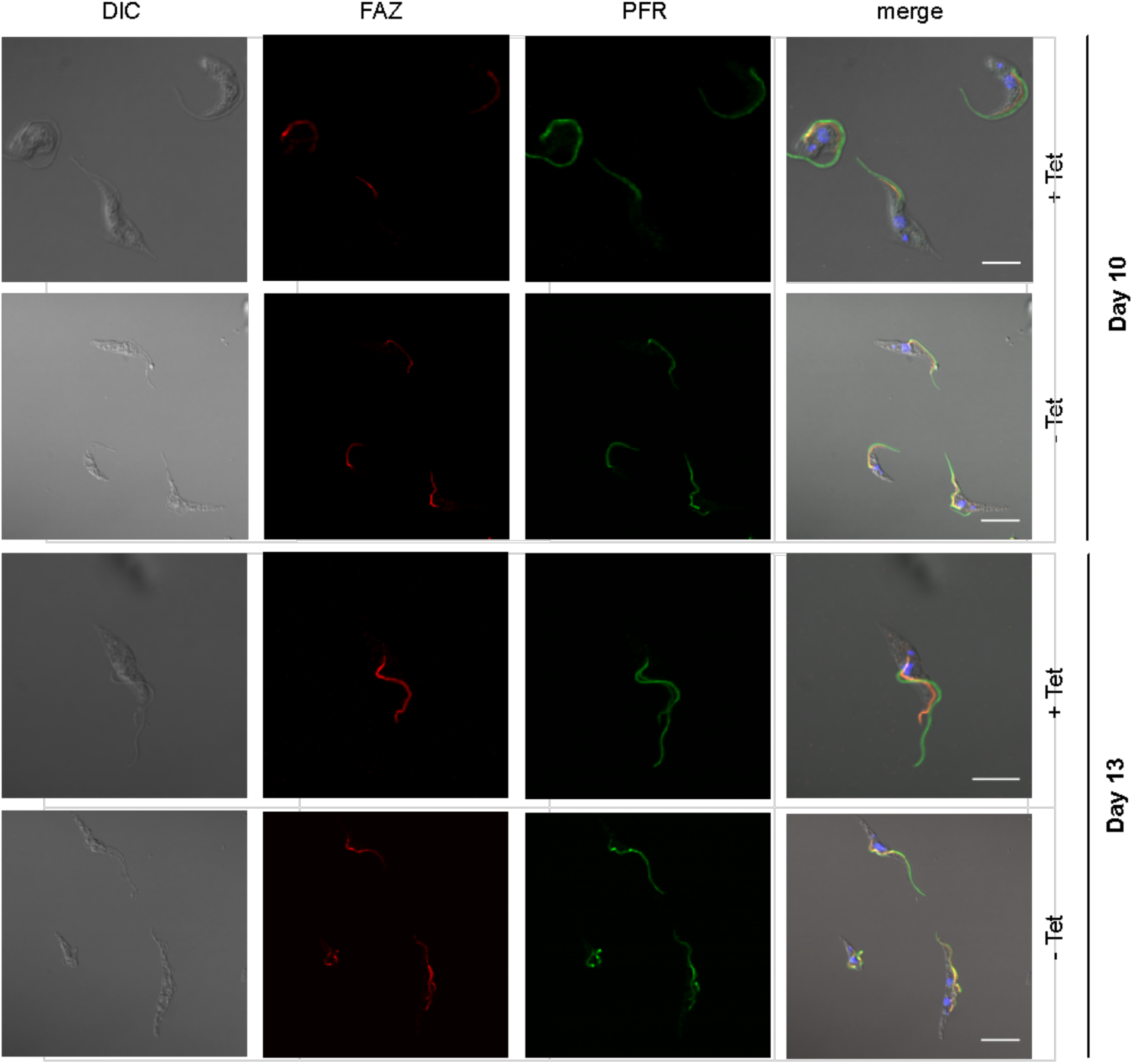
The paraflagellar rod (PFR) and flagellar attachment zone (FAZ) appear normal in PCF *TbFUT1* cKO. Cells were grown in permissive *(+tet*) or non-permissive *(-tet*) conditions and harvested at days 10 and 13. They were fixed, permeabilized and stained as described in the Experimental Procedures. The antibodies were used at the following dilutions: anti-FAZ 1:2, anti-PFR 1:10. Scale bars: 4 μm.

**FIGURE S10.**
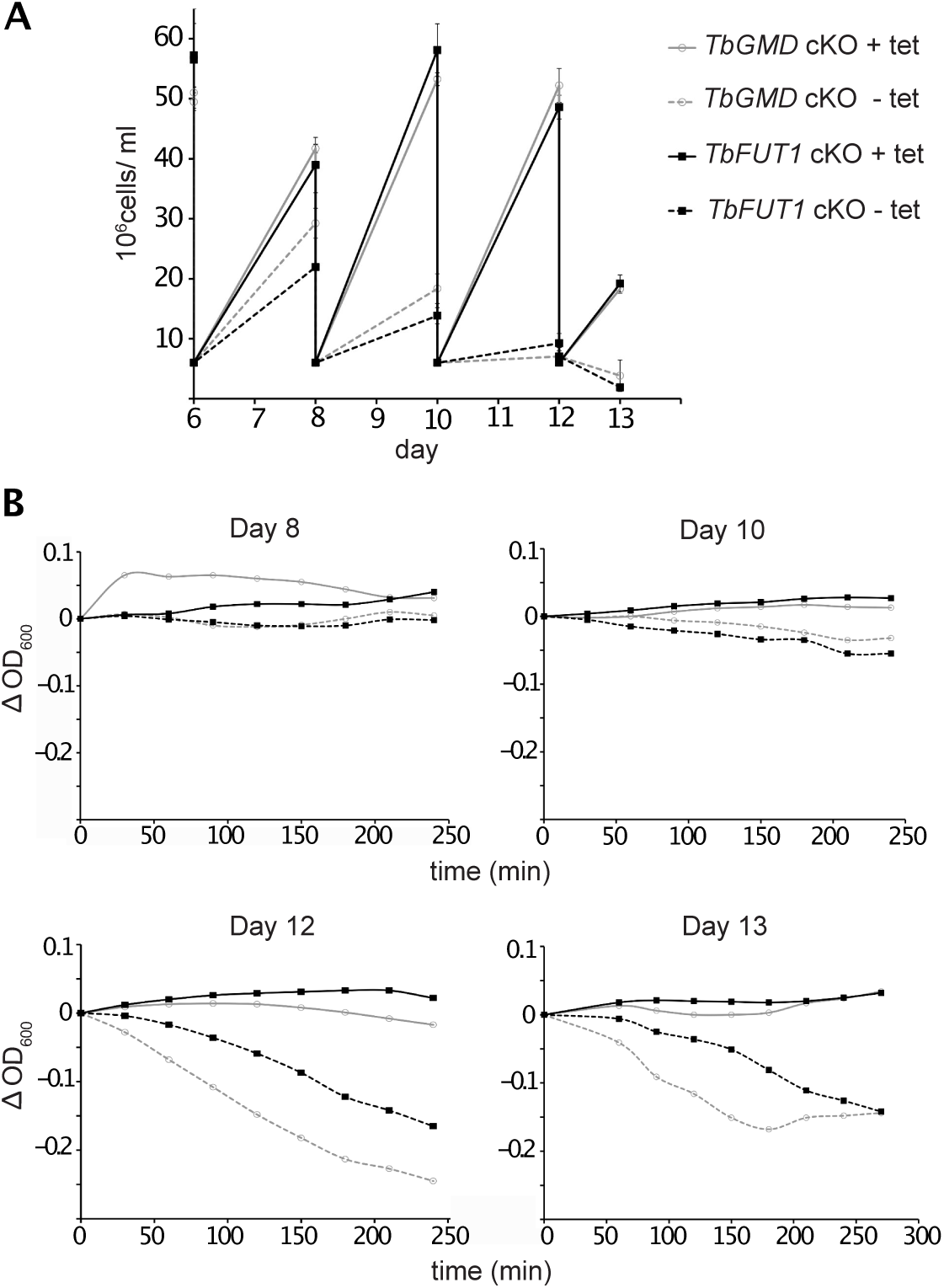
*TbFUT1* and *TbGMD* cKO procyclic form cells show no defect in motility. *A*. Growth curve from day 6 to 13 of the cell lines used in the sedimentation assay. *B*. Sedimentation assays on days 8, 10, 12 and 13. A reduction in motility could be observed in the PCF *TbFUT1* and *TbGMD* cKO cells, grown under non-permissive conditions, only after the start of the growth phenotype (day 12 and onwards), not before. This suggests that the reduction in motility may be an effect of the growth defect and not its cause, as would be expected in a true flagellar attachment mutant (3, 4). *TbGMD* cKO is marked in *gray* while *TbFUT1* cKO is marked in *black*. Cultures grown in non-permissive conditions are marked with *dashed lines*.

**FIGURE S11.**
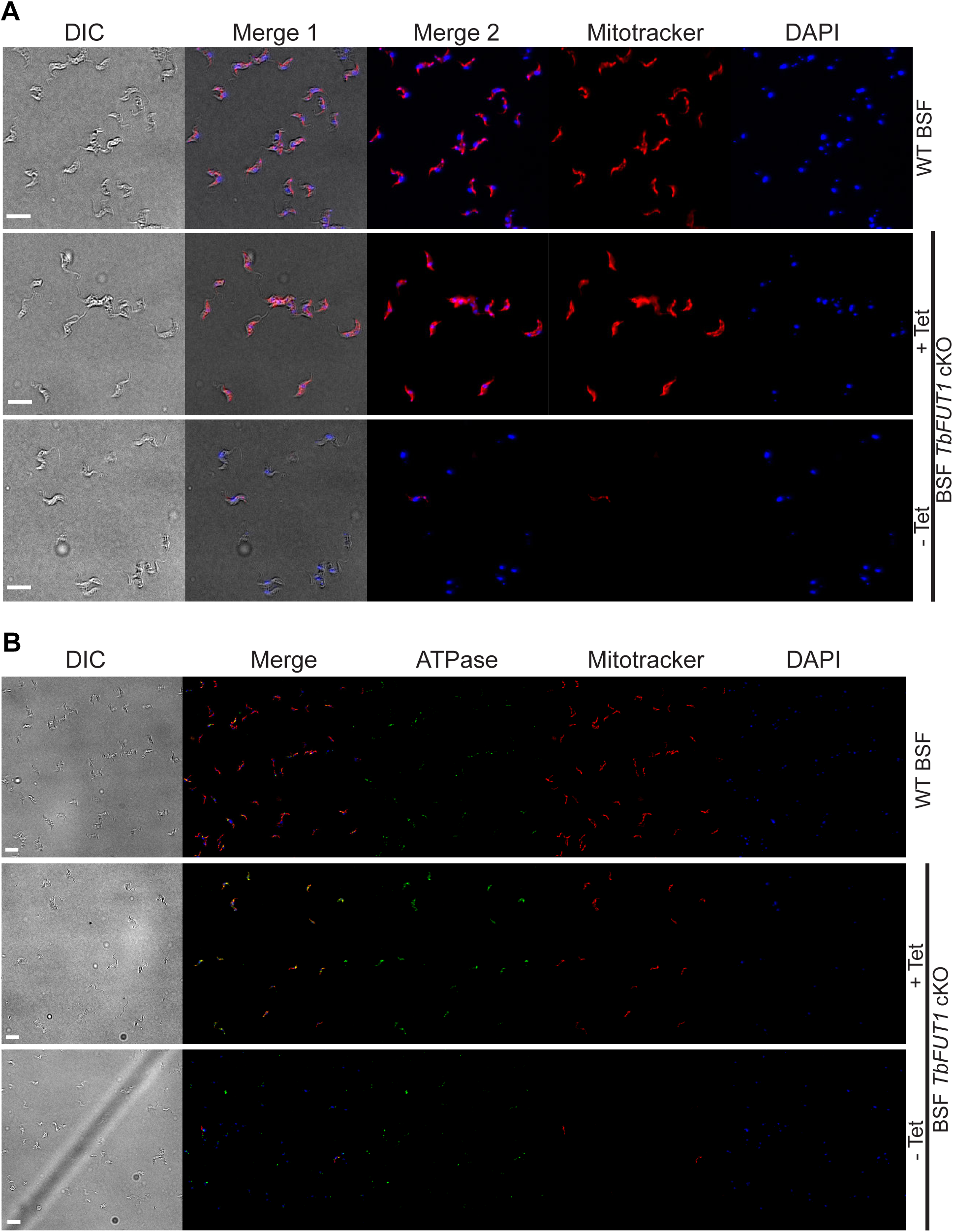
Absence of TbFUT1 disturbs mitochondrial activity. Still viable bloodstream form conditional null mutants cultured for 5 days under non-permissive conditions (-Tet) were tested with MitoTracker^TM^ for mitochondrial activity and co-stained with either DAPI (60x objective) (*A)* or with DAPI and anti-ATPase antibody (*B*) (20x objective). In both cases a strong reduction in Mitotracker^TM^ staining was observed. Scale bars: 10 μm.

